# Src-dependent NM2A tyrosine-phosphorylation regulates actomyosin dynamics

**DOI:** 10.1101/2020.08.12.246090

**Authors:** Cláudia Brito, Francisco S. Mesquita, Daniel S. Osório, Joana Maria Pereira, Neil Billington, James R. Sellers, Ana X. Carvalho, Didier Cabanes, Sandra Sousa

**Author notes:** CB, Cell and Developmental Biology Department, Centre for Genomic Regulation (CRG), Barcelona, Spain; FSM, Global Health Institute, School of Life Sciences, École Polytechnique Fédérale de Lausanne, Lausanne, Switzerland. Shared co-first authorship (Cláudia Brito, Francisco S. Mesquita). Shared second last authorship (Ana X. Carvalho, Didier Cabanes).

## Abstract

Non-muscle myosin 2A (NM2A) is a key cytoskeletal enzyme that along with actin assembles into actomyosin filaments inside cells. NM2A is fundamental in cellular processes requiring force generation such as cell adhesion, motility and cell division, and plays important functions in different stages of development and during the progression of viral and bacterial infections. We previously identified a novel tyrosine phosphorylation on residue 158 (pTyr^158^) in the motor domain of NM2A. This phosphorylation is dependent on Src kinase and is promoted by *Listeria monocytogenes* infection of epithelial cells, however its role is unknown. Here we show that Listeriolysin O (LLO), the pore-forming toxin (PFT) secreted by *L. monocytogenes*, is sufficient to trigger NM2A pTyr^158^ by activating Src, an upstream regulator of actomyosin remodeling. We further address the role of NM2A pTyr^158^ on the organization and dynamics of the actomyosin cytoskeleton and find that, by controlling the activation of the NM2A, the status of the pTyr^158^ alters cytoskeletal organization, dynamics of focal adhesions and cell motility. *In vitro*, we observe that non-phosphorylatable and phospho-mimetic versions of NM2A at Tyr^158^ display motor and ATPase activities similar to the wild-type NM2A, which indicates that the phenotype of these mutants in cells is independent of their ability to translocate actin filaments. Importantly, we find the regulation of this phosphorylation site to be of physiological relevance in *Caenorhabditis elegans*, in particular in response to intoxication by a PFT and to heat shock. We conclude that the control of the phosphorylation status at NM2A Tyr^158^ is a conserved trait that contributes to the regulation of actomyosin dynamics and the ability of cells to respond to bacterial infection. We propose Src-dependent NM2A pTyr^158^ as a novel layer of regulation of the actomyosin cytoskeleton.

## Introduction

The non-muscle myosin 2A (NM2A) is a major cytoskeletal enzyme that plays a central role in the contractility of the actin cytoskeleton [1]. Each NM2A molecule is a hexameric complex composed of two non-muscle myosin heavy chains (NMHC2As) and two pairs of light chains: the regulatory (RLCs) and essential (ELCs) light chains [1, 2]. NMHC2A folds into three different domains: the conserved head/motor domain comprising the sites for ATP hydrolysis and actin binding; the neck domain that interacts with the light chains, stabilizing the protein and regulating NM2A activity; and the tail containing a coiled-coil region responsible for heavy chain homodimerization and NMII filament formation [2, 3]. Phosphorylation of the RLCs at threonine 18 and serine 19 regulates NM2A enzymatic activity and filament formation [4–7]. The inactive form of NM2A with unphosphorylated RLCs folds into a compact structure in which motor heads and tails directly interact [2, 8, 9]. This autoinhibited molecule does not form filaments, interacts only weakly with actin and freely diffuses in cells [10–13]. Upon phosphorylation of the RLC, active NM2A molecules undergo antiparallel self-association to form bipolar filaments [10, 14], which interact with actin filaments and power actomyosin network contraction by converting chemical energy into mechanical force *via* ATP hydrolysis [15]. NM2A filament assembly and their subcellular distribution are additionally regulated by a specific phosphorylation on RLC tyrosine 155 in migrating cells [16] and on serine and threonine residues at the NMHC2A tail domain [17, 18].

NM2A is broadly required for processes dependent on force generation and remodeling of the actomyosin cytoskeleton [1, 19]. Defects on NM2A function are associated with a multitude of human diseases [20–22], including various viral and bacterial infections [23]. Upon cellular infection by the human pathogen *Listeria monocytogenes* (*Lm*), Src kinase is activated [24] and NMHC2A is phosphorylated in the conserved tyrosine 158 (Tyr^158^), located in its motor domain [25]. Phosphorylation of Tyr^158^ (pTyr^158^) occurs in a Src-dependent manner and restricts *Lm* cellular invasion [25]. NM2A is also involved in host cellular responses to plasma membrane (PM) damage induced by bacterial pore-forming toxins (PFTs) [26–29]. Upon intoxication, NM2A coordinates actomyosin cytoskeleton reorganization, cellular blebbing and lysosomal secretion, which are required for PM repair and host survival responses to PFTs [26, 28, 30]. Whether NMHC2A Tyr^158^ is triggered during the actomyosin cytoskeleton reorganization that is required for PM repair is unknown.

Here we investigate if the remodeling of the actomyosin network caused by PFTs [26, 28] correlates with NMHC2A pTyr^158^ triggered by pathogens during cellular infection [25]. We also assess the impact of NMHC2A pTyr^158^ on myosin activity and dynamics, cell adhesion and migration. We find that, although the phosphorylation status of Tyr^158^ does not interfere with the motor and ATPase properties of NM2A *in vitro,* the Src-dependent regulation of pTyr^158^ is essential to control the organization of the actomyosin cytoskeleton, assembly/disassembly of focal adhesions and cell motility. *Caenorhabditis elegans* CRISPR/Cas-9-modified animals expressing phospho-mimetic or non-phosphorylatable versions of NMY-2, an essential non-muscle myosin-2 in this system, are sterile and more sensitive to stress and to intoxication, respectively. These data emphasize the phosphorylation of NMHC2A in Tyr158 as a conserved mechanism with physiological relevance at the single cell and organism level.

## Results

### 1. LLO intoxication induces Src kinase activation

Given that *Lm* infection triggers the Src-dependent pTyr^158^ of NMHC2A [25] and that its PFT Listeriolysin O (LLO) promotes the remodeling of the NM2A cytoskeleton [26, 27], we investigated if LLO by itself is sufficient to trigger NMHC2A pTyr and to activate Src kinase. Using an anti-pTyr antibody, total pTyr proteins were immunoprecipitated (IP) from lysates of HeLa cells in control conditions (non-intoxicated, NI) or after incubation with purified LLO. While the levels of NMHC2A in whole cell lysates (WCL) were comparable in NI and LLO-intoxicated samples, NMHC2A was significantly enriched in pTyr IP fractions from LLO-intoxicated cells (Fig 1A, B). This, together with our previous data [25], indicates that LLO is sufficient to trigger NMHC2A pTyr. To assess if LLO-induced NMHC2A pTyr correlates with the activation of Src kinase, we evaluated the levels of activated Src in non-intoxicated (NI) and LLO-intoxicated HeLa cells. Full activation of Src requires both dephosphorylation of its tyrosine 530 (Tyr^530^) and phosphorylation of its tyrosine 419 (Tyr^419^) [31, 32], specifically detected by immunoblot. Both the levels of Src phosphorylated on Tyr^419^ (Src pTyr^419^) and dephosphorylated on Tyr^530^ increased upon LLO-intoxication of HeLa cells (Fig 1C-F). To confirm that LLO is sufficient to trigger Src activation, we examined the distribution of active Src in single cells using the specific FRET-based ECFP/YPet Src biosensor [33]. HeLa cells expressing the biosensor were imaged and LLO was added after 10 min. The ratio of CFP/YPet signals, indicating active/inactive Src, remained constant throughout time in NI conditions (Fig 1G, H and Movie M1). Upon LLO intoxication the CFP/YPet ratio increased and remained high over time (Fig 1G, H), reflecting global and lasting Src activation in LLO-intoxicated cells. High CFP/YPet values were detected in plasma membrane (PM) blebs (Fig 1G, arrows), where Src kinase mainly acts and the cell is responding to LLO-induced PM damage. Concurrently, confocal microscopy analysis of NI and LLO-intoxicated HeLa cells showed that active Src mainly localized at the perinuclear region in NI cells, but upon LLO intoxication, it was also detected at the cell cortex within the previously described PFT-induced NM2A-enriched cortical bundles [26, 28] (Fig 1I). This indicates that Src activity might be required at the cell periphery to regulate the cytoskeleton remodeling triggered by LLO-induced PM damage. In addition, these data indicate that LLO is sufficient to activate Src, which in turn may control NMHC2A pTyr as it occurs during *Listeria* infection [25].

**Figure 1.**
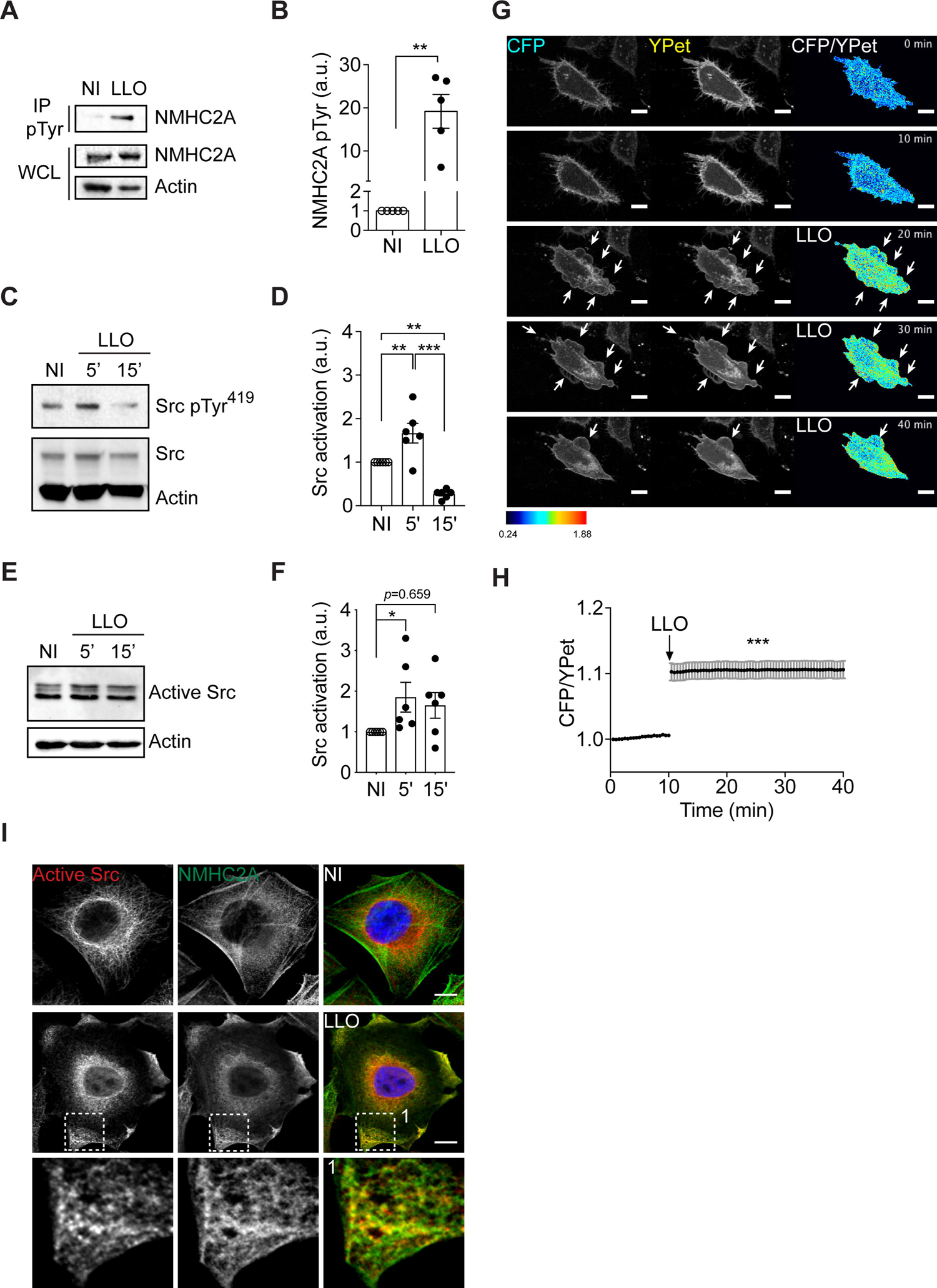
LLO promotes NMHC2A pTyr and Src activation in HeLa cells. **(A, B)** Levels of NMHC2A measured by immunoblots on whole-cell lysates (WCL) and IP fractions of pTyr proteins (IP pTyr) from non-intoxicated (NI) or LLO-intoxicated (LLO; 0.5 nM, 15 min) HeLa cells. Actin was used as loading control. **(B)** The level of NMHC2A in the IP pTyr fraction was quantified and normalized to that detected in the WCL. Each dot corresponds to an independent experiment. Data correspond to mean ± SEM (n=5); *p*-value was calculated using two-tailed unpaired Student’s *t*-test, \*\**p* < 0.01. **(C-F)** Levels of active Src assessed by immunoblots on whole-cell lysates of HeLa cells non-intoxicated (NI) or intoxicated with 0.5 nM LLO for 5 and 15 min. Actin was used as loading control. **(C)** Immunoblot showing the levels of Tyr^419^-phosphorylated Src (Src pTyr^419^) and total Src. **(D)** Quantification of Src pTyr^419^ signals normalized to the levels of total Src. Each dot corresponds to an independent experiment. Values are the mean ± SEM (*n* = 6); *p*-values were calculated using one-way ANOVA with Tukey’s *post hoc* analysis, \*\**p* < 0.01 and *** *p* < 0.001. **(E)** Immunoblot showing the levels of active Src through detection of Src non-phosphorylated at Tyr^530^**. (F)** Quantification of active Src (Tyr^530^ non-phospho Src) signals normalized to the actin levels. Each dot corresponds to a single independent experiment. Values are the mean ± SEM (*n* = 6); *p*-values were calculated using two-tailed unpaired Student’s *t*-test, \**p* < 0.05. **(G)** Sequential frames of time-lapse confocal microscopy FRET analysis of NI or LLO-intoxicated HeLa cells expressing ECFP/YPet-based Src biosensor. LLO was added to the culture medium 20 s before acquisition. Arrows indicate LLO-induced plasma membrane (PM) blebbing sites. Scale bar, 10 µm. **(H)** Kinetics of donor/acceptor fluorescence emission ratio (ECFP/YPet) of the Src biosensor in NI and LLO-intoxicated cells. Values correspond to the mean ± SEM (*n* > 30); *p*-values were calculated using one-way ANOVA with Dunnett’s *post hoc* analysis, \*\*\**p* < 0.001. **(I)** Confocal microscopy images of NI or LLO-intoxicated (0.5 nM, 15 min) HeLa cells, immunolabeled for active Src (red) and NMHC2A (green) and stained with DAPI (blue). Insets (1) show LLO-induced cortical NMHC2A structures enriched in active Src. Scale bar, 10 µm.

### 2. Src kinase coordinates actomyosin cytoskeleton remodeling

Next, we investigated whether the activation of Src kinase interferes with LLO-induced NM2A-enriched cortical bundles (hereafter designated NM2A structures), which are assembled during repair and survival responses to PFTs [26]. LLO-treated HeLa cells pre-incubated with the Src pharmacological inhibitor Dasatinib (Dasa) [34] or expressing low levels of Src (through specific shRNAs) were immunostained for NMHC2A and analyzed by confocal microscopy as compared to controls (Fig 2A). The percentage of cells displaying LLO-induced NM2A-enriched cortical bundles was not affected by perturbation of either Src activity or expression (Fig 2B). However, in Src-deficient conditions, NM2A structures were smaller and seemed to be more enriched in NM2A when compared to controls (Fig 2A). In addition, while the majority of control cells displayed 1 or 2 NM2A structures, the percentage of cells displaying at least 3 structures was significantly increased in Src-deficient cells (Fig 2C, D). These data indicate that, while dispensable for the assembly of LLO-induced NM2A cortical structures, Src activity is necessary to coordinate the remodeling of the cytoskeleton in response to intoxication. We thus hypothesized that Src activity is critical for efficient cellular response to PM damage. We measured, by flow cytometry, LLO-induced PM damage following propidium iodide (PI) incorporation in NI or LLO-intoxicated HeLa cells, under control or Src-deficient conditions. As expected, the percentage of PI-positive cells significantly increased after LLO-intoxication (Fig 2E). This increase was more pronounced in cells with impaired Src activity (Dasa treated) or with low Src expression (shSrc) (Fig 2E), indicating that active Src limits LLO-induced PM damage. These results demonstrate that Src activity regulates the actomyosin cytoskeleton remodeling and contributes to PM repair upon LLO-intoxication.

**Figure 2.**
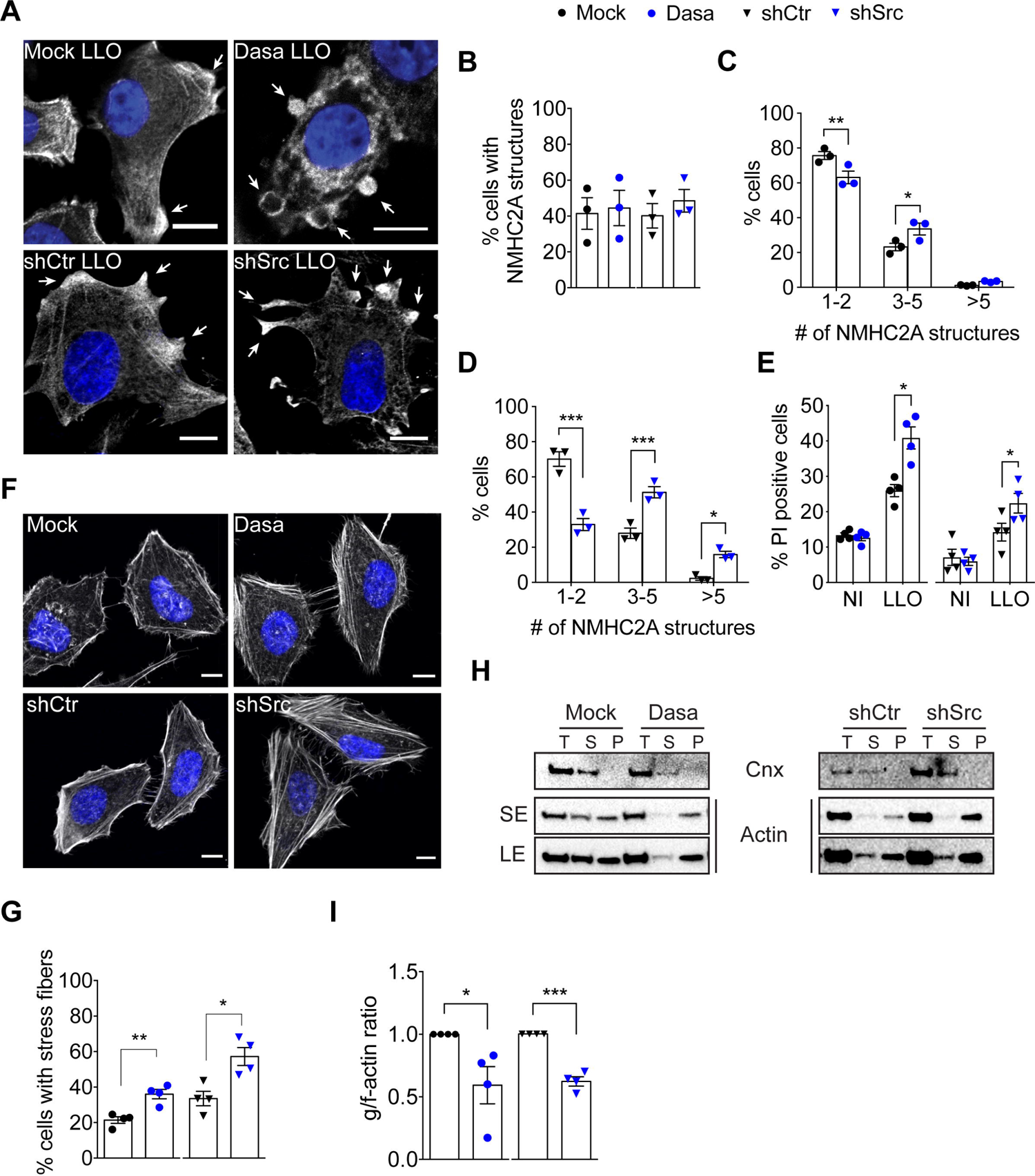
Src kinase orchestrates the response to LLO-intoxication by controlling actomyosin cytoskeleton dynamics in HeLa cells. **(A)** Confocal microscopy images of LLO-intoxicated (0.5 nM, 15 min) HeLa cells in control (mock and shCtr) and Src-impaired (Dasatinib-treated and shSrc) conditions. Cells were immunolabeled for NMHC2A (greyscale) and stained with DAPI (blue). White arrows show NMHC2A cortical structures. Scale bar, 10 µm**. (B)** Percentage of cells displaying NMHC2A structures in mock, Dasatinib-treated (Dasa), shCtr and shSrc HeLa cells, intoxicated as in (A). Values are the mean ± SEM (*n* = 3). **(C, D)** Percentage of LLO-intoxicated cells showing 1-2, 3-5 or >5 NMHC2A structures, under the conditions as in (A). Values are the mean ± SEM (*n* = 3); *p*-values were calculated using two-way ANOVA with Sidak’s *post hoc* analysis, * *p* < 0.05, ** *p* < 0.01, *** *p* < 0.001. (**C)** Shows data from cells under Src chemical inhibition (Dasa). **(D)** Shows data from cells in which Src expression was downregulated through shRNAs. **(E)** Percentage of PI-positive HeLa cells in non-intoxicated (NI) conditions or LLO-intoxicated as in (A), measured by flow cytometry analysis. Values are the mean ± SEM (*n* = 4); *p*-values were calculated using one-way ANOVA with Dunnett’s *post hoc* analysis, \**p* < 0.05. **(F)** Confocal microscopy images of control (mock and shCtr) and Src-impaired (Dasatinib-treated and shSrc) HeLa cells stained with phalloidin for actin (greyscale) and with DAPI (blue). Scale bar, 10 µm. **(G)** Percentage of cells with stress fibers quantified from images similar to those shown in (F). Values are the mean ± SEM (*n* = 4); *p*-values were calculated using two-tailed unpaired Student’s *t*-test, \**p* < 0.05, \*\**p* < 0.01. **(H)** Immunoblots showing the levels of globular (g)- and filamentous (f)-actin in HeLa cells treated as in (F). g- and f-actin from the total cell lysates (T) were separated by ultracentrifugation. g-actin was recovered from supernatant fractions (S) while f-actin was associated with pellet fractions (P). Calnexin was used as loading control (Cnx). Actin levels are shown with both short exposure (SE) and long exposure (LE). **(I)** Quantification of g-/f-actin ratio obtained from immunoblot signals. Values are the mean ± SEM (*n* = 4); *p*-values were calculated using two-tailed unpaired Student’s *t*-test, \**p* < 0.05, \*\*\**p* < 0.001. In **(B, C, D, E, G and I)** each dot or triangle corresponds to an independent experiment.

We further analyzed the role of Src activity in the overall organization of the actomyosin cytoskeleton in non-intoxicated cells. Cells with reduced Src activity or expression consistently showed more stress fibers than control cells (Fig 2F). The overall percentage of cells displaying stress fibers significantly increased under impaired Src conditions (Fig 2G). To support these findings, we performed actin fractionation assays to measure the levels of globular and filamentous actin (g-actin and f-actin, respectively). Src-impaired cells showed decreased levels of g-actin (in the supernatant fraction, S) and concomitantly, increased levels of f-actin (pellet fraction, P) (Fig 2H). In agreement with increased stress fibers detected by fluorescence microscopy, the g-/f-actin ratio was significantly reduced in Src-impaired cells (Fig 2I). Together, these data indicate that besides being critical in coordinating the cytoskeletal responses to LLO-induced PM damage, Src activity also controls the overall actomyosin cytoskeleton organization in HeLa cells growing in standard culture conditions.

### 3. Src kinase activity drives LLO-induced NMHC2A pTyr

To determine if Src activity is necessary for NMHC2A pTyr mediated by LLO intoxication, we manipulated Src activity by: 1) pharmacological inhibition using Dasatinib (Dasa, Fig 3A, B), 2) downregulation using Src specific shRNAs (shSrc, Fig 3C, D), and 3) overexpression of Src kinase mutants (Fig 3E, F). Levels of NMHC2A were assessed in IP pTyr fractions obtained from LLO-intoxicated cells in control or Src perturbed conditions. In control conditions, LLO consistently induced an enrichment of NMHC2A in the IP pTyr fraction (Fig 3A-F). In contrast, NMHC2A pTyr was reduced upon Dasatinib treatment (Dasa; Fig 3A, B), downregulation of Src kinase expression (shSrc; Fig 3C, D) or ectopic overexpression of a Src kinase dead mutant (KD; Fig 3E, F). Inversely, upon LLO intoxication, the ectopic overexpression of a constitutively active (CA) Src mutant increased NMHC2A pTyr to levels higher than those detected in cells expressing wild-type Src (WT) (Fig 3E, F). Notably, low expression levels of Src CA and Src KD when compared to that of WT Src were sufficient to modulate the levels of NMHC2A pTyr (Fig 3E, F).

**Figure 3.**
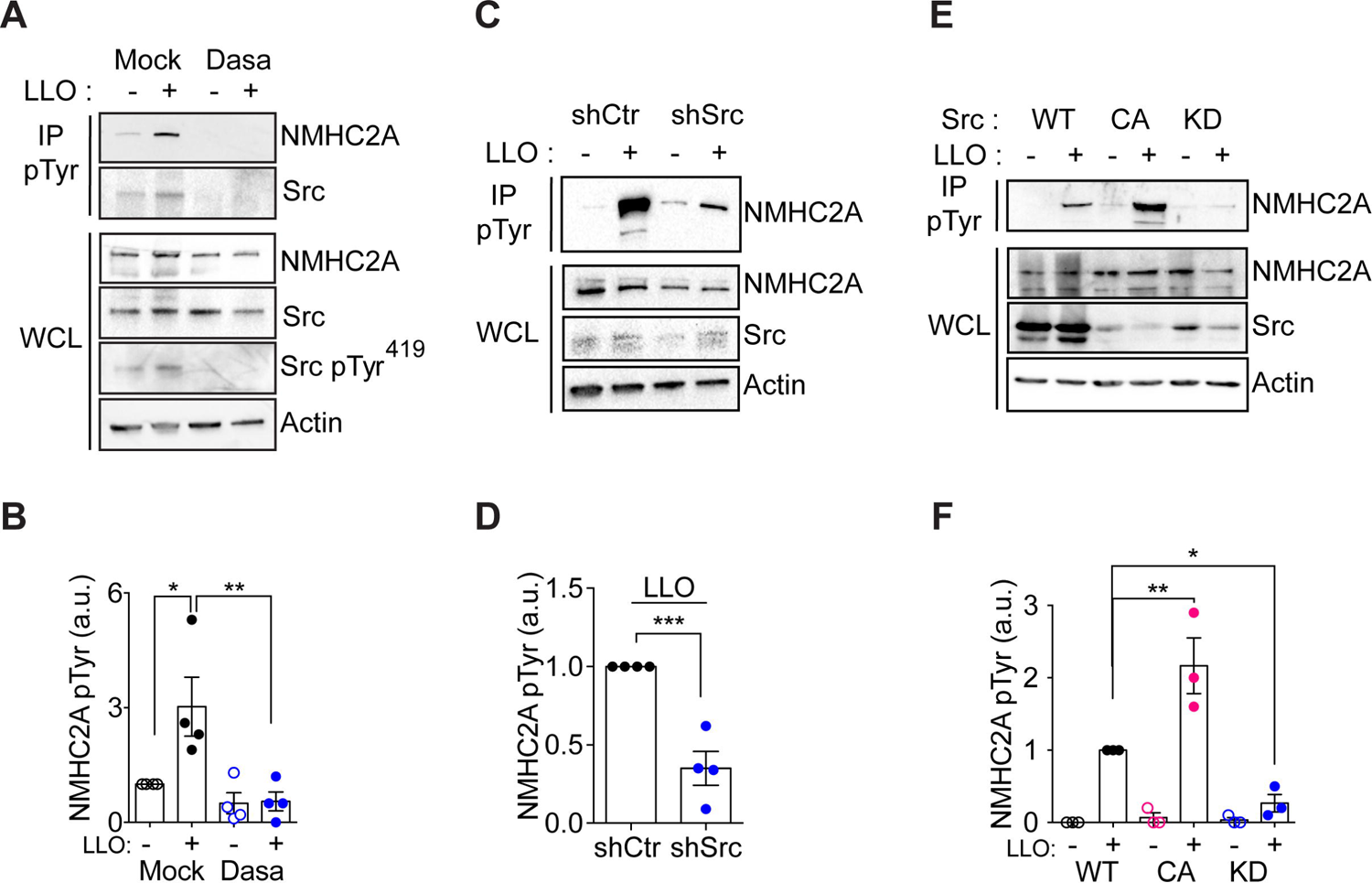
LLO-induced NMHC2A pTyr requires Src activity. Immunoblots for NMHC2A, total Src and Src pTyr^419^ on WCL or IP pTyr fractions of non-intoxicated (-) or LLO-intoxicated (0.5 nM, 15 min) (+) HeLa cells **(A)** in absence (mock) or presence of Dasatinib (300 nM, 1 h) (Dasa); **(C)** expressing control (shCtr) or Src-specific oligos (shSrc) and **(E)** ectopically expressing wild-type Src (WT), constitutively active Src (CA) or kinase dead Src (KD) variants. Actin was used as loading control. **(B, D, F)** Show the quantified levels of NMHC2A in IP fractions normalized to those detected in WCL, under the different conditions. Each dot corresponds to an independent experiment. Values correspond to the mean ± SEM (*n* ≥ 3) and *p*-values were calculated using **(B, F)** one-way ANOVA with Dunnett’s *post hoc* analysis, \**p* < 0.05, \*\**p* < 0.01 or **(D)** two-tailed unpaired Student’s *t*-test, \*\*\**p* < 0.001.

Altogether, these results show that LLO intoxication triggers NMHC2A pTyr specifically through Src kinase activation and induces cortical localization of Src active form, where it is necessary to control actomyosin cytoskeleton remodeling and to assist PM damage repair.

### 4. NMHC2A pTyr does not affect kinetic and structural properties of NM2A in vitro

Tyr^158^ is localized in the proximity of NM2A ATPase domain [25] (Fig S1A) and is conserved throughout evolution (Fig S1B). We thereby investigated if phosphorylation of Tyr^158^ perturbs NM2A ATPase activity or actin-binding, by determining the mechanical and kinetic parameters of human NMHC2A heavy meromyosin (HMM) fragments, which do not form filaments but retain biological activity. We used HMM fragments derived from wild-type NMHC2A (NM2A-GFP-Flag^HMM-WT^), a non-phosphorylatable mutant in which Tyr^158^ was substituted by a phenylalanine (Phe, NM2A-GFP-Flag^HMM-Y158F^) and a mutant mimicking a permanent phosphorylation in which Tyr^158^ was replaced by a glutamate (Glu, NM2A-GFP-Flag^HMM-Y158E^). Such HMM fragments were co-expressed and co-purified with the RLCs and ELCs from Sf9 insect cells (Fig S2A). Purified molecules were analyzed by electron microscopy and no apparent conformational changes were detected among the different HMM variants (NM2A-GFP-Flag^HMM-WT/Y158F/Y158E^, Fig S2B), suggesting that the substitution of Tyr^158^ by a Phe or a Glu do not affect the overall HMM structure. The enzymatic activity of the NM2A-GFP-Flag^HMM^ variants was measured in a steady-state ATPase assay. Although the mutant molecules appeared to hydrolyze ATP at a slightly higher rate than the WT NM2A-GFP-Flag^HMM^ (Fig 4A), this difference was not significant and the K*_ATPase_* (concentration of actin required for one half V_max_) and V_max_ values obtained were similar (Fig 4B), and comparable to those previously described for WT NM2A [35]. This indicates that the pTyr status of the NMHC2A does not affect the ATPase activity of NM2A. The binding affinity of NM2A-GFP-Flag^HMM^ variants to actin was then assessed by co-sedimentation assays in the absence or presence of ATP. As expected, actin alone was pelleted by ultracentrifugation, whereas NM2A-GFP-Flag^HMM^ variants remained in the supernatant in the absence of actin (Fig 4C). NM2A-GFP-Flag^HMM^ molecules co-sedimented with actin in the absence of the ATP but mainly remained in supernatants in the presence of 1 mM ATP (Fig 4C). The fraction of each NM2A-GFP-Flag^HMM^ variant bound to actin in the presence of ATP was comparable in all conditions (Fig 4D). Corroborating these data, electron microscopy analysis of mixtures of actin and the different NM2A-GFP-Flag^HMM^ variants in the presence or absence of ATP showed similar behaviors (Fig S2C). Finally, we assessed if NMHC2A Tyr^158^ is involved in the velocity with which actin filaments slide on NM2A-GFP-Flag^HMM^. We performed *in vitro* actin-gliding assays using coverslip-immobilized NM2A-GFP-Flag^HMM^. All the NM2A-GFP-Flag^HMM^ variants translocated actin at similar rates (Fig 4E and Movie M2), comparable to those previously reported for human NM2A [36].

**Figure 4.**
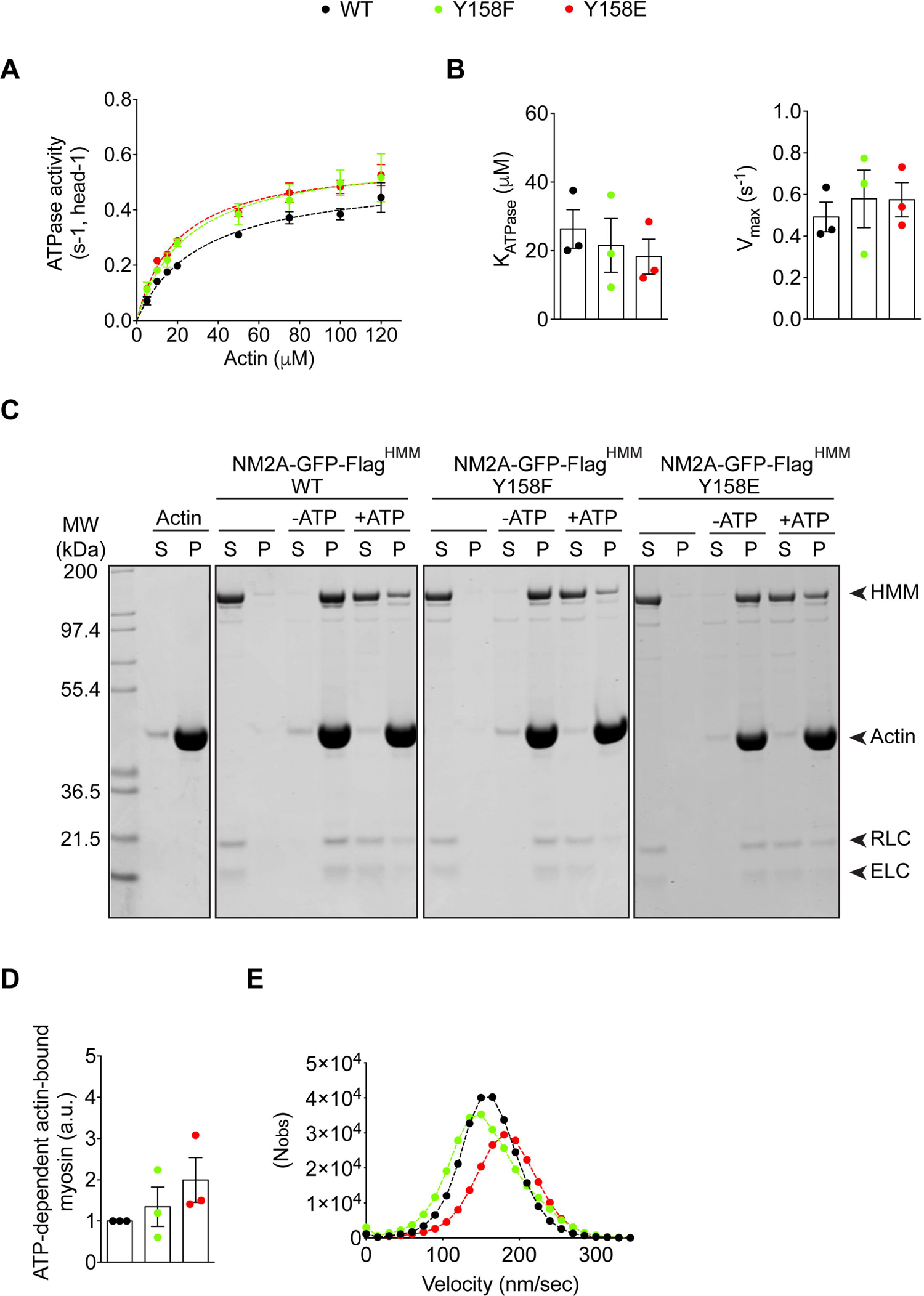
NMHC2A pTyr does not affect myosin 2A (NM2A) mechanical or kinetic properties. **(A)** Actin-activated Mg-ATPase activity of heavy meromyosin (HMM) NM2A-GFP-Flag^HMM^-WT, -Y158F and -Y158E determined by the conversion of NADH into NAD^+^. Three independent preparations of each NM2A-GFP-Flag^HMM^ variant were used in at least 3 independent experiments. Data sets were fitted to a hyperbolic equation to determine the kinetic constants, K_ATPase_ and V_max_. **(B)** Plot of the K_ATPase_ and V_max_ values. Data represent the mean ± SEM from at least 6 independent experiments (per actin concentration) and using the 3 different preparations of NM2A-GFP-Flag^HMM^ variants. Each dot represents the value obtained for a single preparation. Values are the mean ± SEM (*n* = 3). **(C)** Representative actin co-sedimentation assay of NM2A-GFP-Flag^HMM^ WT, -Y158F and -Y158E. Supernatants (S) and pellets (P) obtained from ultracentrifugation of mixtures of either NM2A-GFP-Flag^HMM^ -WT, - Y158F or -Y158E with or without 10 µM actin and 1mM ATP, are shown by Coomassie blue staining. Actin alone is also shown. Regulatory light chain (RLC); Essential light chain (ELC). **(D)** Quantification of co-sedimentation assays showing myosin fraction bound to actin (pellet) in presence of ATP. Each dot corresponds to a single preparation of each NM2A-GFP-Flag^HMM^ variant. Values are the mean ± SEM (*n* = 3). **(E)** Gaussian distribution of the velocity of actin filaments moving on top of either NM2A-GFP-Flag^HMM^-WT, -Y158F or -Y158E. Values were obtained from at least 5 independent motility assays (Movie M2) performed on 3 independent NM2A-GFP-Flag^HMM^ preparations. 500 to 1300 filaments were quantified for each condition and per independent experiment.

These results demonstrate that the substitution of the Tyr^158^ by a Phe or Glu does not significantly affect the *in vitro* biochemical properties of NM2A, indicating that NMHC2A pTyr^158^ does not directly interfere with the regulation of NM2A ATPase activity or the ability to bind and translocate f-actin *in vitro*.

### 5. The status of NMHC2A pTyr contributes to the organization of the actomyosin cytoskeleton

In cells, the NMHC2A pTyr^158^ status may nevertheless control events such as cytoskeleton organization through NM2A filament assembly or spatiotemporal regulation. We thus hypothesized that NMHC2A pTyr^158^ could contribute to Src-induced actomyosin remodeling observed in standard conditions and during intoxication. To address this, we tested the impact of NMHC2A pTyr^158^ in the overall organization of the actomyosin cytoskeleton. HeLa cells were transfected with constructs of GFP-tagged wild-type NMHC2A (GFP-NMHC2A-WT) or phospho-mimetic/non-phosphorylatable mutants (GFP-NMHC2A-Y158E and GFP-NMHC2A-Y158F, respectively). The percentage of transfected cells (Fig S3A) and the mean GFP signal intensity (Fig S3B) were similar for the three constructs, as assessed by flow cytometry.

GFP-NMHC2A-Y158E aggregated in 40% of the cells while GFP-NMHC2A-WT was detected in aggregates in only 20% of the cells (Fig 5A, B), suggesting that GFP-NMHC2A-Y158E may have folding and/or assembly defects and may trigger toxic effects in cells. This observation together with the fact that phospho-mimetic mutations do not always mimic the presumed phosphorylations [37], led us to pursue our study with only the non-phosphorylatable mutant (GFP-NMHC2A-Y158F).

**Figure 5.**
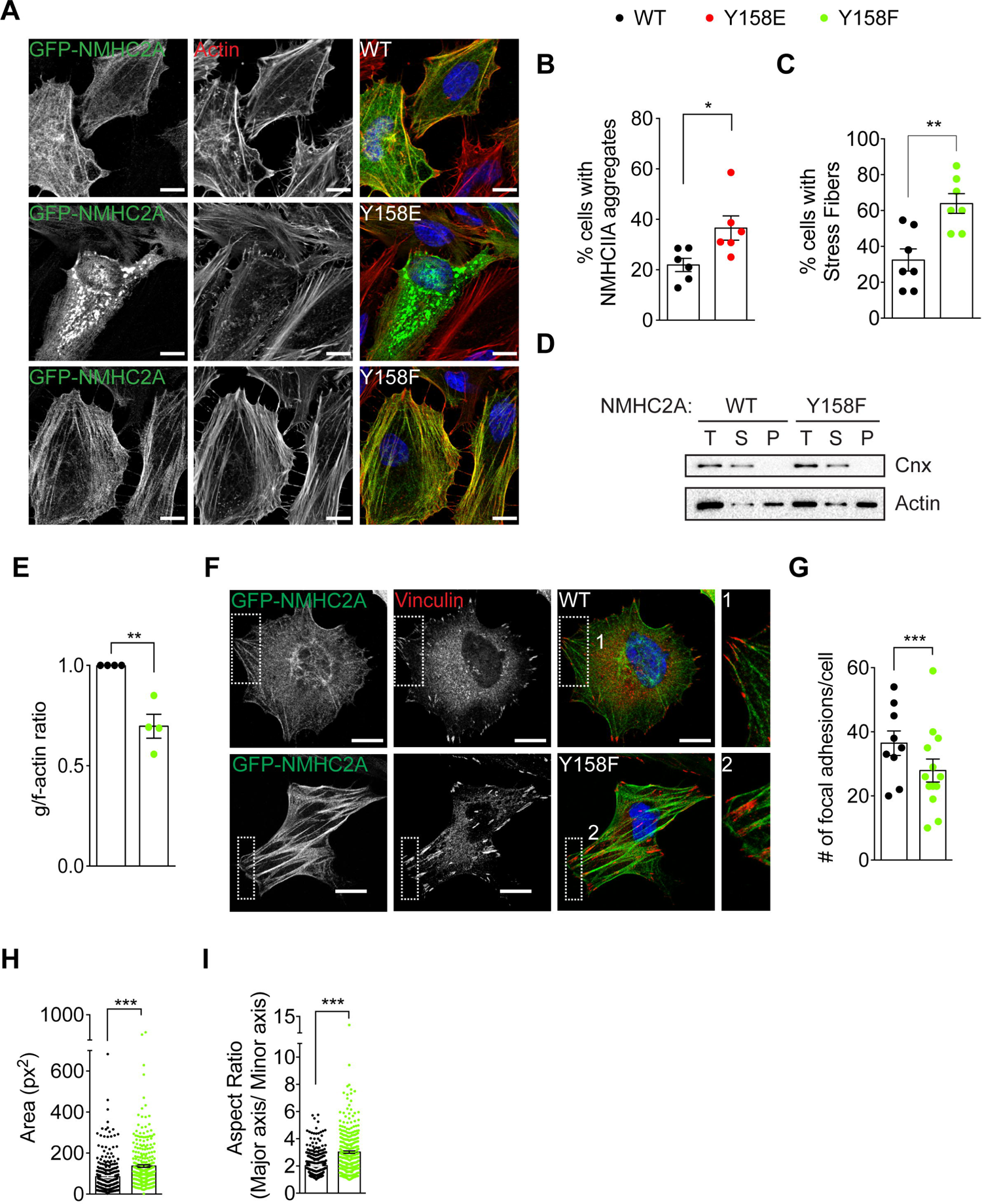
NMHC2A pTyr regulates stress fiber and focal adhesion formation in HeLa cells. **(A)** Representative confocal microscopy images of HeLa cells ectopically expressing either GFP-NMHC2A-WT, -Y158E or -Y158F (green) and stained with phalloidin for actin (red) and DAPI (blue). Scale bar, 10 µm. **(B, C)** Scoring of the percentage of cells displaying **(B)** NMHC2A aggregation or **(C)** stress fibers in HeLa cells ectopically expressing different NMHC2A variants. Each dot corresponds to an independent experiment. Values are the mean ± SEM (*n* ≥ 6); *p*-values were calculated using two-tailed unpaired Student’s *t*-test, \**p* < 0.05, \*\**p* < 0.01. **(D)** Immunoblots showing the levels of globular (g)- and filamentous (f)-actin in HeLa cells ectopically expressing NMHC2A WT or Y158F variants. g- and f-actin from the total cell lysates (T) were separated by ultracentrifugation. g-actin was recovered from supernatant fractions (S) while f-actin was associated with pellet fractions (P). Calnexin was used as loading control (Cnx). **(E)** Immunoblots quantification showing the g/f-actin ratio. Each dot corresponds to an independent experiment. Values are the mean ± SEM (*n* = 4); *p*-value was calculated using two-tailed unpaired Student’s *t*-test, \*\**p* < 0.01. **(F)** Representative confocal microscopy images of HeLa cells ectopically expressing GFP-NMHC2A-WT or GFP-NMHC2A-Y158F (green), immunolabeled for vinculin (red) and stained with DAPI (blue). Insets show focal adhesion morphology. Scale bar, 10 µm. **(G)** Quantification of the number of focal adhesions per cell, from images similar to those shown in F. Each dot corresponds to the mean number of focal adhesions per cell obtained from 3 independent experiments. Values are the mean ± SEM; *p*-value was calculated using two-tailed unpaired Student’s *t*-test, \**p* < 0.05. **(H, I)** Quantification of the focal adhesion **(H)** area and **(I)** aspect ratio, from images similar to those shown in F. Each dot corresponds to a single focal adhesion. Values are the mean ± SEM (*n* > 140); *p*-values were calculated using two-tailed unpaired Student’s *t*-test, \*\*\**p* < 0.001.

The majority of cells expressing GFP-NMHC2A-Y158F were found enriched in stress fibers. They displayed at least 50% of their area covered by stress fibers (Fig 5A, C) organized in an anisotropic orientation (Fig S4). GFP-NMHC2A-WT expressing cells only occasionally showed stress fibers covering more than 50% of their area, which also displayed anisotropic orientation (Fig 5A, C, S4). Actin fractionation assays on total lysates of transfected cells confirmed that the amount of f-actin in the pellet fraction was higher in GFP-NMHC2A-Y158F than in GFP-NMHC2A-WT expressing cells (Fig 5D) and thus, the g-/f-actin ratio was significantly decreased in the former (Fig 5E). Together these results reveal that cells expressing the non-phosphorylatable GFP-NMHC2A-Y158F accumulate stress fibers and f-actin and phenocopy the results obtained in cells with reduced Src activity (Fig 2F-I). NMHC2A pTyr through Src is thus important to regulate cell cytoskeleton organization.

Given that stress fibers and focal adhesions are structurally and functionally interdependent and that focal adhesions maturation requires NMII [38, 39], we investigated whether focal adhesions would also be affected by the NMHC2A pTyr status. Cells overexpressing GFP-NMHC2A-WT or -Y158F were fixed and immunolabelled for vinculin, a marker for focal adhesions [40]. The number, size and shape of focal adhesions were analyzed by confocal microscopy. Although the number of focal adhesions per cell was slightly decreased in GFP-NMHC2A-Y158F expressing cells (Fig 5F, G), they were larger and more elongated, displaying increased area and aspect ratio (Fig 5F, H, I), which may suggest that in these cells focal adhesions are under increased tension. These data suggest that pTyr status of NMHC2A not only regulates stress fiber formation but also interferes with focal adhesions and their associated tension. Overall our data show the importance of NMHC2A pTyr^158^ in the overall cytoskeletal organization and suggest its involvement in cell adhesion and migration.

### 6. NMHC2A pTyr interferes with cell size and controls cell motility

To test if the status of NMHC2A pTyr^158^ affects cell shape parameters and cell migration, we compared the area and aspect ratio of cells expressing GFP-NMHC2A-WT or -Y158F. Expression of GFP-NMHC2A-Y158F led to slightly increased cell area (Fig 6A). Yet, aspect ratio values were similar in both conditions (Fig 6B).

**Figure 6.**
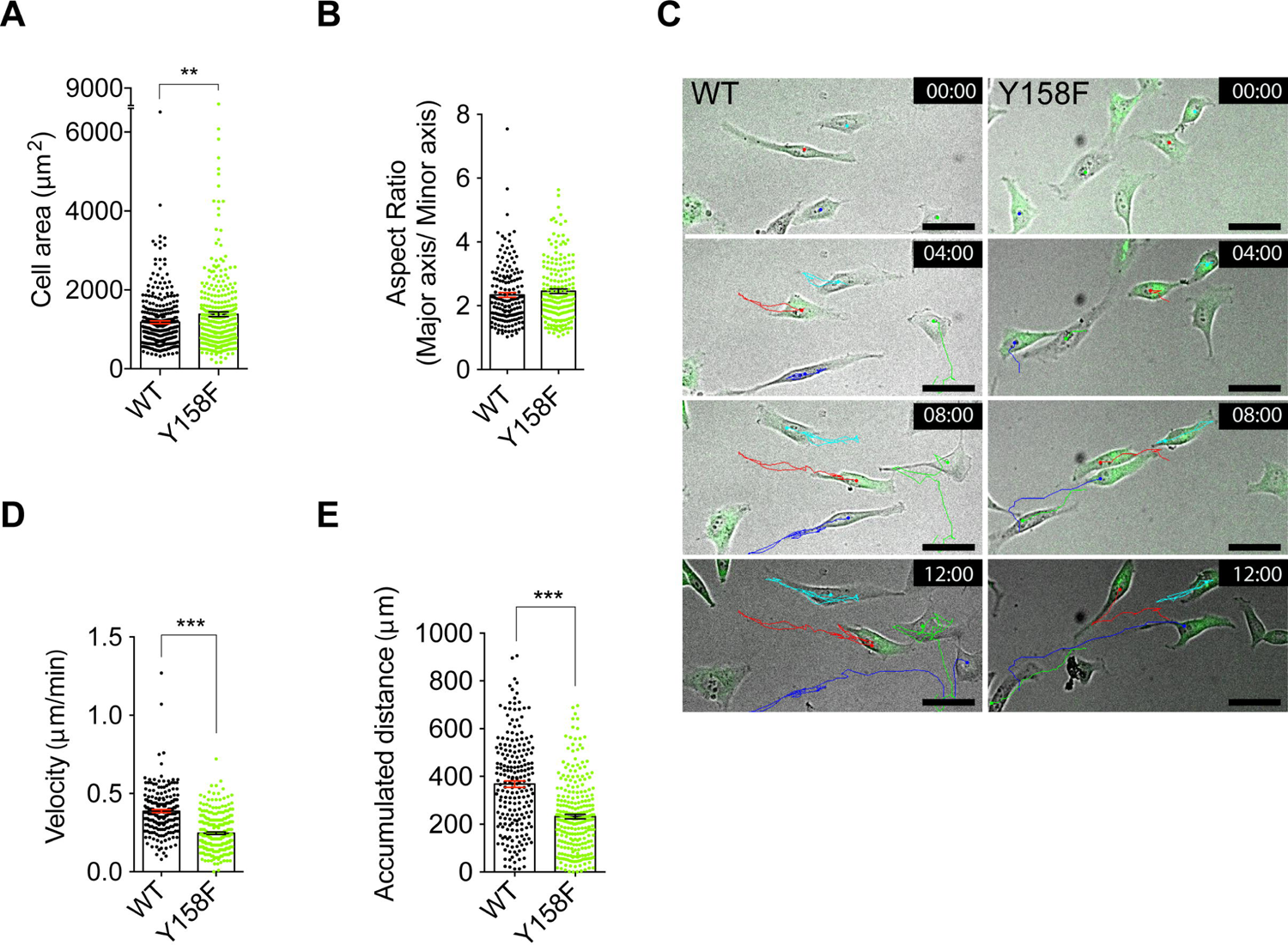
NMHC2A pTyr status alters cell size and motility. **(A, B)** Quantification of cell shape parameters of HeLa cells ectopically expressing the GFP-NMHC2A-WT or -Y158F: **(A)** area and **(B)** aspect ratio. Cells were stained with phalloidin for actin and quantifications were performed using Fiji™. Each dot represents a single cell. Values are the mean ± SEM (*n* > 160); *p*-values were calculated using two-tailed unpaired Student’s *t*-test, \*\*\**p* < 0.001. **(C)** Sequential frames of the time-lapse microscopy Movie M3, from 0 until 10 h 30 min. HeLa cells ectopically expressing GFP-NMHC2A-WT or GFP-NMHC2A-Y158F are shown. The movement of individual cells over time was analyzed using the Fiji™ Manual Tracking plugin and is represented by color lines. Scale bar, 50 μ **(D, E)** Quantification of cell motility parameters using the Manual Tracking plugin of Fiji™ **(D)** velocity and **(E)** accumulated distance in HeLa cells as in A. The data was obtained from movies similar to Movie M3. Each dot represents a single cell. Values are the mean ± SEM (*n* > 200); *p*-value was calculated using two-tailed unpaired Student’s *t*-test, **p < 0.001.

To further evaluate the consequences of the cytoskeletal alterations induced by NMHC2A pTyr^158^, we monitored the motility of HeLa cells using fluorescent time-lapse microscopy (Fig 6C and Movie M3). Cell-tracking analysis of time-lapse videos revealed that cells expressing GFP-NMHC2A-Y158F migrated with reduced velocity (Fig 6D) and spanned shorter distances (Fig 6E), as compared to cells expressing GFP-NMHC2A-WT. These results are in line with the elongated focal adhesions and increased stress fibers we observed, indicating that cells are possibly under more tension if the non-phosphorylatable NMHC2A-Y158F is expressed. The NMHC2A pTyr^158^ status thus modulates cell motility, most likely through the control of the actomyosin cytoskeleton organization.

### 7. NMHC2A pTyr regulates NM2A activation and dynamics

We next examined whether pTyr^158^ could affect NM2A turnover. To investigate this, we performed Fluorescence Recovery After Photobleaching (FRAP) in HeLa cells overexpressing either the GFP-NMHC2A-WT or -Y158F. The GFP signal of NM2A was photobleached in a single region per cell where NMHC2A filaments were detected, the recovery of the signal was followed over time and the data were analyzed using easyFRAP-web on-line platform [41]. The GFP signal within the photobleached region in cells overexpressing GFP-NMHC2A-Y158F recovered faster than that in cells overexpressing GFP-NMHC2A-WT (Fig 7A, B). In addition, while the GFP signal reached a plateau in GFP-NMHC2A-Y158F, such signal continuously increased in GFP-NMHC2A-WT cells without reaching a plateau during the time of analysis (Fig 7B). The mobile fraction, corresponding to the fraction of molecules able to turnover, and the half time required for signal recovery (T-half) were obtained from easyFRAP-web mean curve fitting tool [41]. Calculated mobile fractions were similar in both conditions (Fig 7C). However, the half-time for signal recovery was significantly shorter in cells expressing GFP-NMHC2A-Y158F as compared to those expressing GFP-NMHC2A-WT (Fig 7D), indicating that the expression of non-phosphorylatable NMHC2A-Y158F allows for faster NM2A turnover.

**Figure 7.**
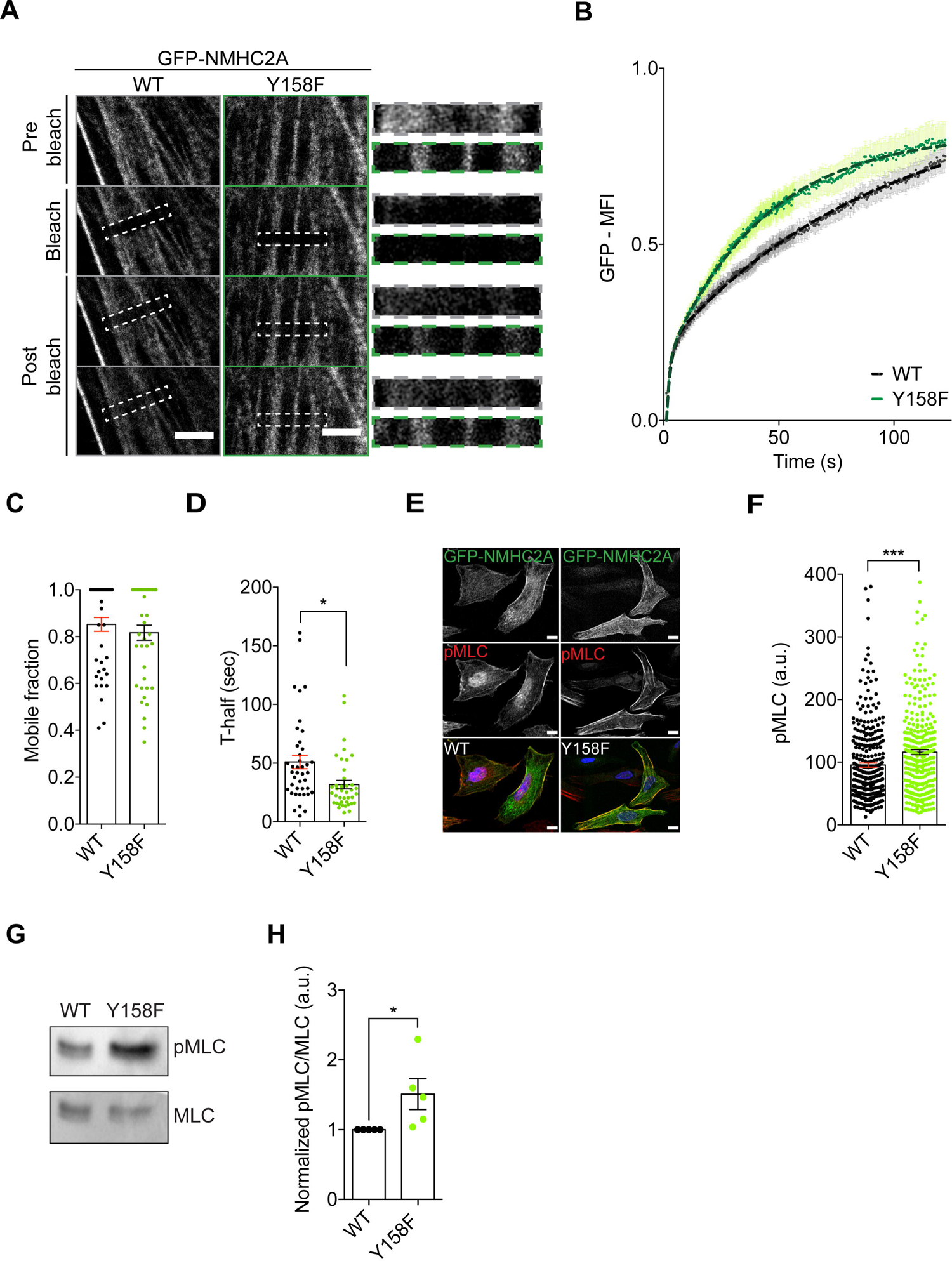
NMHC2A pTyr^158^ controls the dynamic assembly of bipolar filaments in HeLa cells. **(A)** Representative confocal microscopy time-lapse images of Fluorescence Recovery After Photobleaching (FRAP) experiments, in HeLa cells ectopically overexpressing either the GFP-NMHC2A-WT or -Y158F. Insets of the bleached region in dashed lines. Scale bar, 5 μm **(B)** Plot of the normalized mean fluorescence intensity recovery curves obtained for each condition tested (black, WT; green, Y158F). Values are the mean ± SEM (n ≥ 40 cells). **(C, D)** Quantification of the **(C)** mobile fraction and the **(D)** half-maximal recovery time (T-half) obtained from the curve fitting of each individual recovery curve. Each dot represents a single cell. Values are the mean ± SEM (n ≥ 40 cells); *p*-values were calculated using two-tailed unpaired Student’s *t*-test, \*\**p* < 0.01. **(E)** Representative confocal microscopy images of HeLa cells ectopically expressing either the GFP-NMHC2A-WT or -Y158F (green), immunolabeled to detect pMLC (red) and stained with DAPI (blue). Scale bar, 15 µm. **(F)** Levels of pMLC in cells expressing the different NMHC2A variants, quantified using Fiji™. Each dot corresponds to a single cell. Values are the mean ± SEM (*n* > 300); *p*-value was calculated using two-tailed unpaired Student’s *t*-test, \*\*\**p* < 0.001. **(G)** Immunoblot showing the levels of phosphorylated MLC (pMLC) and the total levels of MLC on whole cell lysates of HeLa cells expressing either GFP-NMHC2A-WT or -Y158F. **(H)** Quantification of pMLC immunoblot signal normalized to that of MLC. Each dot corresponds to an independent experiment. Data correspond to mean ± SEM (n=5); *p*-value was calculated using two-tailed unpaired Student’s *t*-test, \**p* < 0.05.

NM2A filament dynamics and self-organization require NM2A contractility, which is in turn regulated by RLC phosphorylation [2, 42]. Thus, we quantified phosphorylated RLC in residues threonine 18 and serine 19 (pMLC) to assess the levels of NM2A activation. In agreement with a faster turnover, we found that cells overexpressing the non-phosphorylatable GFP-NMHC2A-Y158F displayed higher levels of pMLC than those expressing GFP-NMHC2A-WT (Fig 7E-H). In addition, pMLC co-localized with stress fibers induced by the expression of GFP-NMHC2A-Y158F (Fig 7E). These results, together with the *in vitro* biochemical data, suggest that non-phosphorylated Tyr^158^ trigger NM2A activation and increase its dynamics within assembled filaments and not at the single molecule level.

### 8. Deregulated pTyr of NMHC2A homologue impairs toxin and stress responses in C. elegans

Nematodes are naturally susceptible to infection caused by PFT-secreting bacteria such as *Bacillus thuringiensis*, which produces Cry5B, a PFT that reduces *C. elegans* viability [43]. Given that host responses to PFTs and NMII function are conserved in nematodes, we used the *C. elegans* model together with Cry5b intoxication approach to evaluate the role of NMHC2A pTyr^158^ *in vivo*. *C. elegans* NMY-2 is homologous to the human NMHC2A (Fig S5) and NMY-2 functions were linked to Src-mediated signaling. Thus, we introduced point mutations by CRISPR/Cas9 in the *nmy-2* genomic locus replacing the conserved tyrosine 163 (Tyr^163^) by a non-phosphorylatable (Y163F) or a phosphomimetic (Y163E) residue. To test the effects of each mutation in *C. elegans*, we analyzed the brood size (total number of progeny laid) and the embryonic viability (percentage of viable embryos) of each strain. The homozygous strain expressing NMY-2(Y163F) showed normal brood size and embryonic viability. In contrast, homozygous animals expressing NMY-2(Y163E) were sterile and therefore unable to lay embryos (Fig 8A), suggesting that permanent phosphorylation of Tyr^163^ may impact gonad function, or that the presence of this mutation leads to folding defects that impair NMY-2 functions, as suggested by the expression of GFP-NMHC2A-Y158E in HeLa cells.

**Figure 8.**
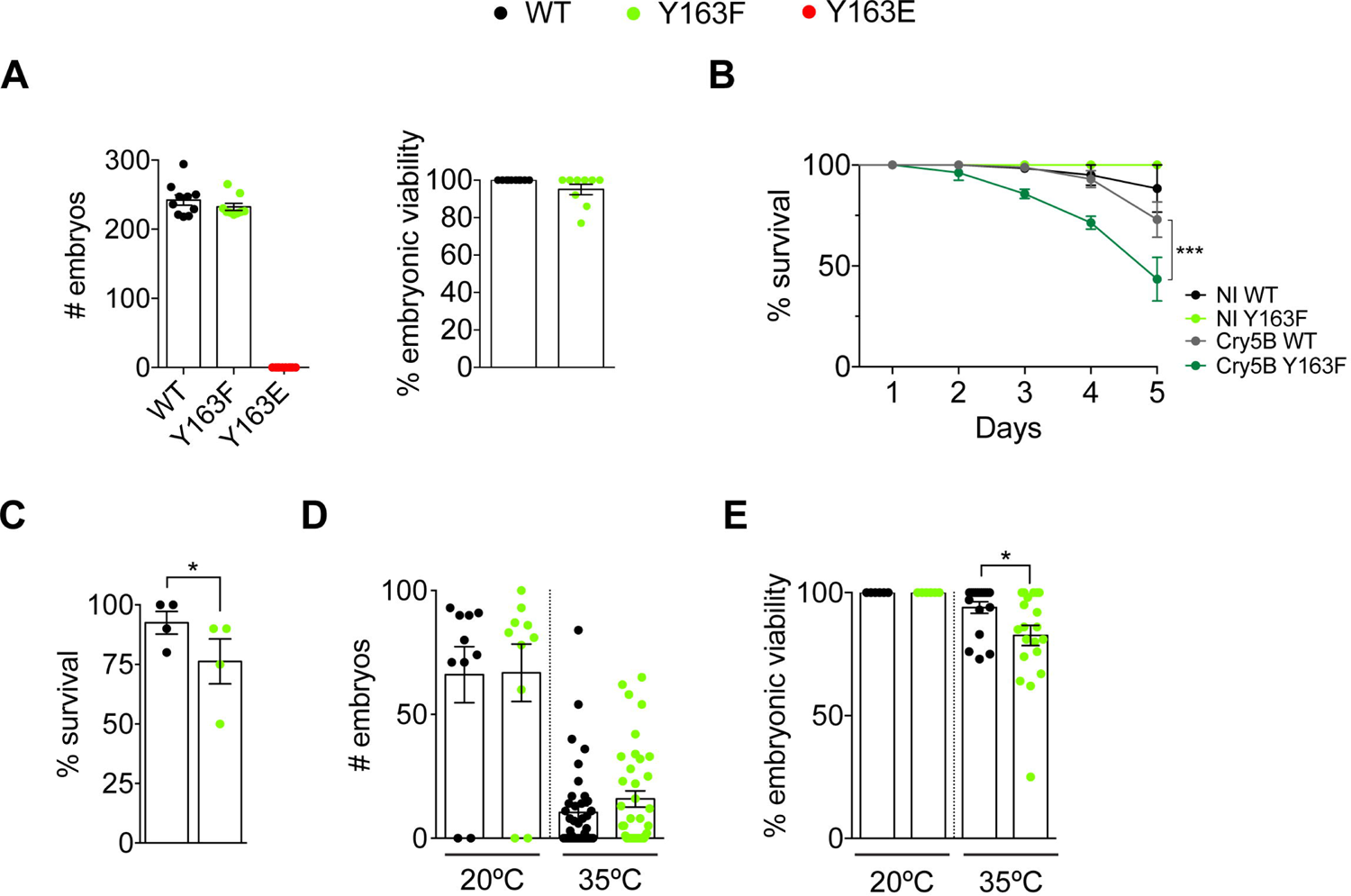
Regulation of NMY2 pTyr on residue 163 is necessary for *C. elegans* survival to intoxication and stress. **(A)** Mean brood size and embryonic viability ± SEM in *C. elegans* expressing wild-type NMY-2 (WT), NMY-2(Y163F) or NMY-2(Y163E). On the left, each dot represents the brood size of a single animal. On the right, each dot represents the embryonic viability (in %) of embryos laid by each animal. **(B)** Mean percentage of survival ± SEM of *wild-type* and *nmy-2(Y163F)* animals fed for 5 days with control *E. coli* (NI WT; NI Y163F) or with *E. coli* expressing the Cry5B toxin (Cry5B WT; Cry5B Y163F). Each curve represents the mean of 3 independent experiments with 30 animals per condition. *p*-values were calculated using two-way ANOVA with Sidak’s *post hoc* analysis, ***p<0.001. **(C)** Mean percentage of survival ± SEM of *wild-type* and *nmy-2*(*Y163F)* animals 24 h post a 1 h heat shock at 35 °C. Each dot represents one independent experiment with 8-10 animals per condition. *p*-values were calculated using two-tailed paired Student’s *t*-test, \**p* < 0.05. **(D)** Mean brood size ± SEM of the animals that survived the 1 h heat shock at 35 °C. Each dot represents the brood size of a single animal. *p*-values were calculated using two-tailed paired Student’s *t*-test, \**p* < 0.05, \*\**p* < 0.01. **(E)** Mean percentage of viability ± SEM of the embryos laid by the animals that survived the 1 h heat shock at 35 °C. Each dot represents the percentage of viable embryos laid by a single animal. *p*-values were calculated using two-tailed paired Student’s *t*-test, \**p* < 0.05, \*\*\**p* < 0.001.

As the animals expressing NMY-2(Y163E) could not be propagated in the homozygous state, we focused our analysis on *wild-type* and *nmy-2(Y163F)* strains. We evaluated their susceptibility to the *B. thuringiensis* PFT, Cry5b. Animals were fed with control or Cry5b-expressing *E. coli* starting at L4 stage and the number of survivors was quantified daily, over a period of 5 days. While wild-type non-intoxicated worms survived and appeared healthy over time, those fed with Cry5b-expressing *E. coli* showed decreased survival (Fig 8B). Animals expressing NMY-2(Y163F) showed normal survival when non-intoxicated but were significantly more susceptible to intoxication than those expressing wild-type NMY-2 (Fig 8B). These results suggest that controlled regulation of NMY-2 phosphorylation at Tyr^163^ is necessary for the response to Cry5b intoxication in *C. elegans*.

To test the possibility that NMY-2 phosphorylation at Tyr^163^ could also be important for the response to other stresses, we assessed its role in response to heat shock. L4 larvae were subjected to heat shock for 1 hour at 35°C and animal survival was assessed 24 hours later. Animals expressing NMY-2(Y163F) showed increased susceptibility to the heat shock (Fig 8C). The number of embryos laid by heat-shocked *wild-type* and *nmy-2(Y163F)* animals was similar (Fig 8D), but embryo viability was lower for heat-shocked *nmy-2*(Y163F) progenitors (Fig 8E).

Altogether, these data demonstrate that in *C. elegans* the regulation of NMY-2 Tyr^163^ phosphorylation is important for survival responses to different stresses, including heat shock and PFT intoxication.

## Discussion

pTyr controls cell signaling and regulates a variety of biological processes, including the reorganization of the cytoskeleton [44, 45]. We previously reported an uncharacterized pTyr event on NMHC2A, triggered by bacterial infections [25]. Infection by *L. monocytogenes* induced Src-dependent NMHC2A phosphorylation on Tyr^158^, a residue that locates close to the ATP-binding pocket of NMHC2A and was predicted to be accessible for phosphorylation [46]. Here we showed that LLO, the *L. monocytogenes* secreted PFT and major virulence factor, is sufficient to trigger NMHC2A pTyr. We also found that the regulation of NMHC2A pTyr^158^ contributes to the cellular organization of the actomyosin cytoskeleton, without interfering with NMHC2A motor or enzymatic activities. In *C. elegans*, the control of this conserved phosphorylation proved to be important for organism survival under intoxication and heat stress conditions.

LLO activation of phosphotyrosine signaling is driven by both pore-dependent and independent mechanisms [47–49]. While Ca^2+^ influx through the LLO pore was proposed as the key event for LLO-induced signaling [50], LLO-mediated clustering of lipid rafts also contributes to tyrosine kinase activation and intracellular signaling responses [49, 51]. Our data demonstrate that during LLO intoxication, activated Src generates a protective response of the host against bacterial-induced PM damage. Paradoxically, Src activity is required for bacterial invasion of host cells [24]. Src has thus a dual role during infection, and its activation might benefit both the host and the pathogen.

Src activity controls cortical tension in NMII-dependent processes and regulates retraction of PM blebs upon osmotic and non-osmotic stress in a MLCK- and/or ROCK-dependent manner [52–56]. PFT-induced PM blebs are extruded or retracted in a process mediated by NM2A and are associated with NM2A-enriched cortical structures [26–28]. We found active Src at sites of LLO-induced PM blebbing, suggesting that Src activation locally modulates the remodeling of the actomyosin cytoskeleton assisting the formation and retraction of PM blebs, which might sustain PM repair mechanisms. High Src activity reduces cell adhesion by triggering stress fiber disassembly [57–59], which correlates with enhanced migration, invasion and tumor metastization [60, 61]. In turn, low Src activity promotes stress fiber assembly associated with cell stiffening [60], and reduced cell migration by interfering with focal adhesions [62]. In line with this, Src-deficient fibroblasts show reinforced interactions between integrins and the actomyosin cytoskeleton [63], also limiting migration. Strikingly, our data show that cells overexpressing a non-phosphorylatable version of NM2A molecule on Tyr^158^ behave similarly to cells with impaired Src activity, suggesting that Src-dependent phosphorylation of NM2A Tyr^158^ is likely to contribute to Src-mediated cell processes.

While the assembly of ventral stress fibers and focal adhesions depend on tension powered by NM2A contractility [64–67], other studies reported that tension and myosin activity had little effect on both stress fibers and focal adhesions assembly [68–70]. Our data favor the involvement of NM2A-mediated mechanical tension in the assembly of stress fibers and their associated focal adhesions. NM2A-Y158F-expressing cells display increased levels of Thr18/Ser19 phosphorylated myosin regulatory light chain and showed faster turnover of NM2A, which might be associated with increased contractility [42]. Larger and more elongated focal adhesions, together with increased stress fibers detected in NM2A-Y158F-expressing cells, suggest an altered contraction status that is associated with increased tension, stronger adhesion to the substratum and increased cell stiffness [60, 71, 72]. The phosphorylation status of NMHC2A Tyr^158^ could therefore regulate the assembly/disassembly of focal adhesions and stress fibers, thus providing a novel mechanism to modulate cytoskeletal organization. Phosphorylation events in the regulatory light chain and the heavy chain tail domain respectively regulate NM2A activity (reviewed in [2]) and the spatial localization [16] and the dynamics of NM2A filaments [73, 74]. To our knowledge pTyr^158^ is the only phosphorylation occurring in the motor domain of NMHC2A that contributes to regulating the dynamics of actomyosin cytoskeleton. Previous studies proposed that the formation of heterotypic filaments [75, 76] and the assembly of NMII supramolecular structures named NMII stacks [42, 77] may modulate actomyosin dynamics likely to accomplish specific functions in particular spatiotemporal contexts [78, 79]. It is possible that pTyr158 regulate the assembly of such heterotypic filaments and/or NMII stacks.

The expression of the phosphomimetic NMY2(Y163E) in *C. elegans* rendered the animals sterile, indicating that this mutation does not sustain gonad function. This agrees with the aggregation phenotype detected in HeLa cells overexpressing NMHC2A-Y158E and may be due to a toxic effect. The expression of motor-dead NMY2 mutants in *C. elegans* leads to sterility in adult homozygous animals [80]. However, the motor activity of the NM2A(Y158E) is not affected *in vitro*, suggesting that the sterility phenotype due to expression of NMY2(Y163E) is independent of its myosin motor activity. NMHC2A pTyr^158^ might be critical for signaling and/or cytoskeletal organization upon stress or environmental assault, however constitutive pTyr^158^ appears to be detrimental. NMY-2 activity and Src-mediated signaling are important to control an asymmetric division of the *C. elegans* embryo that dictates the individualization of the endoderm and mesoderm [81–83]. Activated Src and phosphotyrosine signaling accumulate at the *C. elegans* embryo EMS/P2 cell boundaries to ensure proper orientation of the EMS cell division axis, in a process that was proposed to be NMY-2 dependent [81, 83]. In addition, Src plays a key role in directing the distal tip cell migration in *C. elegans* during gonad morphogenesis [82]. NMY-2 functions and Src-mediated signaling appear thus intimately linked in *C. elegans*, however the mechanisms through which they regulate each other remain unknown.

We thus propose a new conserved mechanism for the regulation of the actomyosin cytoskeleton through the control of NM2A pTyr^158^. This adds a new layer of complexity in the control of actomyosin dynamics with major impact on cell adhesion to substrate, cell migration, animal development and response to bacterial infection. Our work also reveals the pTyr^158^ as a new event protecting the host cells from bacterial infections, contributing to the current knowledge on the host response to toxins causing PM damage.

## Material and Methods

### Cell lines

HeLa cells (ATCC CCL-2) were grown in DMEM with glucose and L-glutamine (Lonza), supplemented with 10% fetal bovine serum (FBS; Biowest) and maintained at 37 °C in a 5% CO_2_ atmosphere.

### Reagents, toxins, antibodies and dyes

Dasatinib (Santa Cruz Biotechnology) was used at 300 nM in complete medium for 1 hour prior to LLO intoxication. LLO was expressed in *E. coli* BL21(*DE3*) and purified as described in [84]. Intoxications and washes were carried in Hanks’ balanced salt solution (HBSS, Lonza). The following antibodies were used at 1/200 for immunofluorescence microscopy (IF) or 1/1,000 for immunoblotting (IB): rabbit anti-Src pTyr^419^ (#44-660G, Invitrogen), mouse anti-active Src (#AHO0051, Invitrogen), rabbit anti-Src (#sc-18, Santa Cruz), mouse anti-αβ Sigma); mouse anti-phosphotyrosine (#05-321, Millipore); rabbit anti-NMHC2A (#M8064, Sigma); rabbit anti-calnexin (#AB2301, Millipore), rabbit anti-vinculin (#700062, Fisher Scientific), rabbit anti-pMLC (Thr18/Ser19) (#3674, Cell Signaling), rabbit anti-NMY-2 (against residues 945-1368), mouse anti-α-tubulin (#T5168, Sigma). For IF analysis, DNA was stained, 6-diamidino-2-phenylindole dihydrochloride (DAPI; Sigma) and actin with Rhodamine Phalloidin (Thermo Fisher Scientific) at 1/200; secondary antibodies were used at 1/500: goat anti-rabbit Alexa Fluor 488 (Invitrogen), goat anti-rabbit Alexa Fluor 594 (Invitrogen), goat anti-mouse Cy3 (Jackson ImmunoResearch). For IB, goat anti-rabbit or anti-mouse HRP (PARIS) were used at 1/10,000.

### Plasmids

Kras-Src FRET biosensor was a gift from Yingxiao Wang (Addgene plasmid # 78302) [33]. Plasmids allowing the expression of wild-type Src kinase (WT), constitutively active Src (CA) and kinase dead Src (KD) were kindly provided by S. J. Parsons (University of Virginia) [85]. CMV-GFP-NMHCII-A plasmid was a gift from Robert Adelstein (Addgene plasmid # 11347) [86]. pFastBac-NMHC2A^HMM^-GFP-Flag-WT, pFastBac-RLC and -ELC were described in [87]. Insertion of point mutations, Y158F and -Y158E, was achieved by site-directed mutagenesis using QuickChange II Site-directed mutagenesis kit (Agilent) as described in [25].

### Transfections and shRNA lentiviral transductions

HeLa cells seeded on top of glass coverslips in Nunc^TM^ 24-well plates (5×10^4^ cells/well), into Ibitreat μg-dishes (Ibidi) (1×10 cells/well) or Nunc 6-well plates (2.5×10 cells/well) were or 2,5 μ jetPRIME^®^ Polyplus transfection reagent or Lipofectamine 2000^®^ (Thermo Fisher Scientific); according to the manufacturers’ instructions. Protein expression was allowed for 18 to 24 h before cells were processed. shRNA lentiviral transductions were performed as described [25] and the sequences used are available in supplementary material.

### Immunoblot assays

For HeLa cells, protein extracts were recovered in sample buffer (0.25 mM Tris–Cl, pH 6.8; 10% SDS; 50% glycerol; and 5% β-mercaptoethanol), resolved by SDS–PAGE (10% acrylamide) and transferred onto nitrocellulose membranes using a TransBlot Turbo™ system (BioRad), at 0.3 A for 1 h. For *C. elegans*, 60 adult worms/condition were harvested, washed 3 times in M9 containing 0.1% Triton X-100, centrifuged at 750 g, resuspended in sample buffer supplemented with quartz sand (Sigma) corresponding to 1/3 of the final volume. 3 cycles of 5 min boiling at 95°C followed by 5 min vortexing were performed. Samples were resolved by SDS-PAGE (7,5% acrylamide) and transferred onto nitrocellulose membranes (Hybond ECL; GE Healthcare) at 0.22 mA for 2 h at 4°C. Primary and secondary HRP-conjugated antibodies were diluted in TBS–Tween 0.1% (150 mM NaCl; 50 mM Tris–HCl, pH 7.4; and 0.1% Tween) with 5% (m/v) milk. Washes were performed with TBS–Tween 0.2%. Signal was detected using Western Blotting Substrate (Thermo Fisher Scientific) and collected in a ChemiDoc™ XRS+ System with Image Lab™ Software (BioRad).

### Immunofluorescence microscopy

Cells were fixed in 3% paraformaldehyde (15 min), quenched with 20 mM NH_4_Cl (1 h) and permeabilized with 0.1% Triton X-100 (5 min). Coverslips were incubated for 1 h with primary antibodies, washed three times in PBS 1x and incubated 45 min with secondary antibodies and when indicated, Rhodamine Phalloidin and DAPI. Antibodies and dyes were diluted in PBS containing 1% BSA. Coverslips were mounted onto microscope slides with Aqua-Poly/Mount (Polysciences). Images were collected with a confocal laser-scanning microscope (Leica SP5 II) and processed using Fiji™ or Adobe Photoshop software.

### Fluorescence Resonance Energy Transfer (FRET)

HeLa cells were seeded into Ibitreat μ-dishes (Ibidi), transfected to express ECFP/YPet Kras-Src biosensor, maintained in HBSS or HBSS supplemented with LLO (2 nM) at 37°C in 5% CO_2_, and imaged using a confocal laser-scanning microscope (Leica SP8) equipped with a 40x 1.4 NA objective. Non-treated cells were imaged for 10 min prior LLO (2 nM) addition. Image acquisition of LLO-intoxicated cells started 20 s after LLO addition. ECFP/YPet fluorescence m z-steps were acquired every 30 s. Fiji™ was used for image sequence analysis and video assembly. The ECFP/YPet ratio was quantified in the entire surface of more than 30 cells.

### Immunoprecipitation

HeLa cells (6 × 10^6^ cells) treated as indicated were washed with PBS 1x and lysed in RIPA buffer (Santa Cruz). Lysates were recovered after centrifugation at 15,000 x*g* (10 min at 4°C). g) were pre-cleared with protein A-immobilized PureProteome magnetic beads (Millipore) and incubated overnight (4°C) with 1.5 μ antibody (4G10 Millipore). Immune complexes were captured with 50 μ magnetic beads (Millipore) and processed for immunoblotting.

### Flow Cytometry

For flow cytometry, 5×10^6^ cells seeded in six well plates 24 h before treatment, were intoxicated as indicated and washed twice in cold PBS 1x, trypsinized and resuspended in 0.5 ml of cold PBS 1x supplemented with 2% FBS. They were further incubated with 2 μ Iodide (PI, Sigma) 5 min before flow cytometry analysis. At least 30000 events per sample were analyzed.

### g-actin and f-actin separation

HeLa cells (5×10^6^) were seeded in six well plates and either transfected to express GFP-NMHC2A-WT or -Y158F variants; or dasatinib-treated for 1 h. Cells were washed twice with PBS 1x and homogenized 10 min at 37°C in actin stabilization buffer (0.1 M Hepes, pH 6.9, 30% glycerol (v/v), 5% DMSO (v/v), 1 mM MgSO_4_, 1 mM EGTA, 1% Triton X-100 (v/v), 1 mM ATP, complete protease and phosphatase inhibitors). Total protein fractions were ultracentrifuged at 100,000 *g* for 75 min at 37 °C. Supernatants containing g-actin were collected, and pellets containing f-actin resuspended in an equivalent volume of RIPA buffer. Total, supernatant and pellet fractions were processed for immunoblotting.

### NM2A-GFP-Flag^HMM^-WT, -Y158F and -Y158E expression in Sf9 cells

NM2A-GFP-Flag^HMM^ variants were expressed in Sf9 insect cells, following the Bac-to-Bac® Baculovirus Expression System (Life Technologies) as recommended. Briefly, Sf9 cells were co-transfected with the pFastBac-NMHC2A^HMM^-GFP-Flag-WT, or -Y158F, or -Y158E, pFastBac-RLC and -ELC bacmids, and incubated at 27 °C for 96 - 120 h. Supernatants were collected to harvest the viral stock (P1). Sf9 cells in suspension were then infected with P1 (MOI = 1) to obtain a higher viral titer (P2) and incubated at 27 °C with shaking. The process was repeated and the viral titer P3 was used to co-infect Sf9 cells for further expression and purification of the HMMs. Sf9 P3-infected cells were harvested after 72 h and used for purification or stored at −80°C.

### NM2A-GFP-Flag^HMM^-WT, -Y158F and -Y158E purification

Sf9 infected-cell pellets were resuspended in extraction buffer (15 mM MOPS pH 7.3, 200 mM NaCl, 10 mM MgCl_2_, 1 mM EGTA, 3 mM NaN_3,_ 1mM DTT, 0.1 mM PMSF, 5 μ l leupeptin, 2 phosphatase inhibitors tablets) and lysed by sonication. The lysate was incubated with 1mM ATP for 15 min, centrifuged at 19000 rpm for 20 min and incubated with pre-washed anti-Flag-resin (#A2220, Sigma) for 3 h at 4 °C. Flag-resin was washed with high ionic strength buffer (10 mM MOPS pH 7.3, 0.1 mM EGTA, 3 mM NaN_3_, 0.5 M NaCl, 1mM ATP, 5 mM MgCl_2_ and 0.1 mM PMSF), loaded into a column (4 °C) and further washed with 3 column volumes of low ionic strength buffer (10 mM MOPS pH 7.3, 0.1 mM EGTA, 3 mM NaN_3_ and 0.1 mM PMSF). Proteins were eluted with 2 column volumes of Flag-peptide buffer (0.5 mg/ml Flag peptide, 10 mM MOPS pH 7.3, 0.1 mM EGTA, 3 mM NaN_3_, 100 mM NaCl and 0.1 mM PMSF). Eluted fractions were concentrated and dialyzed overnight in dialysis buffer (10 mM MOPS pH 7.3, 0.1 mM EGTA, 3 mM NaN_3_, 0.5 M NaCl and 1mM DTT) at 4°C. Aliquots of the purified proteins were quickly frozen in liquid nitrogen and stored at −80 °C.

### Single molecule Electron Microscopy

For NM2A-GFP-Flag^HMM^ molecules in the absence of actin, samples were diluted to 50 nM HMM in 10 mM MOPS pH 7.0, 50 mM NaCl, 2 mM MgCl_2_, 0.1 mM EGTA. NM2A-GFP-Flag^HMM^ molecules were mixed with F-actin to give final concentrations of 100 nM of HMM and 500 nM of F-actin. Samples were handled as previously described [10]. Briefly, 5 μ sample was applied to a carbon-coated copper grid (pre-exposed to UV light to produce a hydrophilic surface) and stained with 1% uranyl acetate. Data were recorded at room temperature on a JEOL 1200EX II microscope equipped with an AMT XR-60 CCD camera. The Fiji™ FFT bandpass filter (40 – 2 pixels, auto-scaled) was applied to images prior to making figures, in order to enhance the clarity of individual molecules.

### ATPase activity assay

Actin-activated Mg^2+^-ATPase activities were determined by previously described NADH-linked assay [15, 88]. Purified NM2A-GFP-Flag^HMM^ variants were phosphorylated for at least 30 min at room temperature, in adequate buffer conditions (0.2 mM ATP, 0.2 mM CaCl_2_, 10 µg/ml MLCK and 1 µM CaM). Mg^2+^-ATPase activities were determined in 10 mM MOPS, 0.1 mM EGTA, 50 mM NaCl, 2mM MgCl_2_, 3.4 mM ATP, growing actin concentrations (0-120 µM), 40 units/ml lactate dehydrogenase, 200 units/ml pyruvate kinase, 1 mM phosphoenolpyruvate, 0.2 mM NADH and non-phosphorylated or phosphorylated HMMs (0.1 – 0.2 µM). The conversion of NADH into NAD^+^ was determined by measuring the A_340_ (= 6220 M^-1^cm^-1^) every second for 20 min. Data were corrected for the background Mg^2+^-ATPase activity of actin alone. The Mg-ATPase activity of the different NM2A-GFP-Flag^HMM^ variants at 0 µM actin concentration was further subtracted to each data point. To determine the kinetic constants V_max_ and K*_ATPase_*, the experimental data sets were fitted to the Michaelis-Menten mathematical equation using Prism 8.

### Co-sedimentation assay

2 µM of each NM2A-GFP-Flag^HMM^ variant was incubated for 30 min at room temperature with 10 µM f-actin in the absence or presence of 1 mM ATP, in a buffer containing 10 mM MOPS, 0.1 mM EGTA, 100 mM NaCl, 2 mM MgCl_2_, 3 mM NaN_3_, and 1 mM DTT. Samples containing only the NM2A-GFP-Flag^HMM^ variants or filamentous actin were used as sedimentation controls. Samples were centrifuged at 100,000xg for 30 min at 4 °C. Supernatant and pellet fractions were recovered and the loading buffer was added equivalently. Each sample (20 µl) was analyzed on a 4-12% Bis-Tris gel (Life Technologies), stained with Coomassie Blue and scanned on an Odyssey system (Li-Cor Biosciences).

### In vitro motility assay

I*n vitro* motility assays were performed as described [89]. Briefly, 60 μl of NM2A-GFP-Flag^HMM^ molecules (0.2 mg/ml) in motility buffer (MB: 20 mM MOPS, pH 7.3, 0.1 mM EGTA, 2 mM MgCl_2_) with 0.5 M NaCl were trapped in a flow cell. NM2A-GFP-Flag^HMM^ molecules attached to the coverslip were further incubated with 50 μl MB with 0.5 M NaCl and 1 mg/ml BSA, washed in MB with 50 mM NaCl (LS buffer) and incubated for 4 min with 35 μ CaCl_2_, 1mM ATP, 1 μM CaM, 1 nM MLCK and 10 μM of unlabeled actin in LS buffer. Coverslips were washed in LS buffer and incubated with 30 μl of labelled actin filaments (20 nM Rhodamine phalloidin actin in 50 mM DTT) for 35 s. The excess solution was removed and the flow cell was loaded for 1 min with 40 μl of MB with 0.7% methylcellulose, 1mM ATP, 50 mM g/ml glucose oxidase, and 45 μg/ml catalase. The slides were imaged at 25°C, at 5 s intervals for 2 min, on an inverted Nikon Eclipse Ti-E microscope with an H-TIRF module attachment, a CFI60 Apochromat TIRF 100x Oil Immersion Objective Lens (N.A. 1.49, W.D. 0.12 mm, F.O.V 22 mm) and an EMCCD camera (Andor iXon Ultra 888 EMCCD, 1024 × 1024 array, pixel size: 13 μ of the different NM2A-GFP-Flag^HMM^ molecules was quantified using the FAST algorithm described in [90].

### Quantification of immunofluorescence images

*NMHC2A bundles, aggregates, stress fibers, focal adhesions, phospho-regulatory light chain (pMLC), cell area and aspect ratio* The percentage of cells with NM2A bundles and the number of NM2A bundles per cell were quantified in at least 200 cells per sample. Positive cells displayed at least a distinct cortical focus enriched in NM2A, each focus was quantified as a single NM2A bundle. The percentage of cells with stress fibers or NM2A aggregates was quantified in at least 200 cells per sample. Positive cells for stress fibers had approximately 50% (arbitrary) of the total cell area covered by dense actomyosin fibers and positive cells for aggregation displayed at least 2 distinct aggregates larger than 2 µm or one larger than 5 µm. Anisotropy was quantified using the Fiji™ plug-in *FibrilTool* as described in [91] and focal adhesions were quantified as described in [92] using the CLAHE and Log3D Fiji™ plug-ins. The levels of pMLC were quantified using Fiji™ by measuring in each cell the raw integrated density of pMLC normalized by the total cell area in at least 100 cells per sample. The area of the cells and respective aspect ratio were measured using the analysis tool from Fiji™.

### Cell motility analysis

Cells were seeded into Ibitreat μ-dishes (Ibidi), transfected to express the different GFP-NMHC2A variants, maintained in OptiMEM (Gibco) at 37°C and 5% CO_2_, and imaged using a widefield microscope (Leica DMI6000 Time-lapse) equipped with a 20x 0.4 NA objective. The data sets of transmitted light and the fluorescence of GFP-NMHC2A were acquired every 10 min during 16 – 20 h. Fiji™ was used for image sequence analysis and video assembly. The velocity and accumulated distance were analyzed in at least 200 cells/condition using the Manual tracking plug-in from Fiji™.

### Fluorescence Recovery After Photobleaching (FRAP)

HeLa cells were transfected to express either GFP-NMHC2A-WT or -Y158F. Images were acquired 20 h after transfection using a laser scan confocal microscope SP8 (Leica) equipped with a 63x 1.3 NA objective, at 37°C and 5% CO_2_. Five pre-bleach images were acquired each 138 ms, followed by five bleaching scans with 100% intensity on the 488-nm laser line over the region of interest (ROI). Recovery of fluorescence was monitored for 56 s every 138 ms (400 frames), followed by 60 s every 300 ms (200 frames). Mean fluorescence intensities (MFI) were measured on Fiji™ and fluorescence recovery analysis was performed as described in [41], using the easyFRAP-web on-line platform (https://easyfrap.vmnet.upatras.gr/). Briefly, the MFI of the bleached ROI was normalized to non-bleached ROIs. We followed a Full Scale normalization which subtracts background values at each time point and corrects for laser fluctuations, fluorescence loss during photobleaching, differences in starting intensities and loss in total fluorescence. Additionally, the MFI was corrected for differences of the bleaching efficiency following easyFRAP-web on-line platform recommendations [41]. The mobile fraction and the half-maximal recovery time (T-half) values were obtained by curve fitting. Curve fitting was performed using a single and double term exponential equation and obtained R^2^ values served as goodness-of-fit measure. Only cells for which fitted curves showed a R^2^ ≥ 0.8 were considered to quantify the mobile fraction and T-half.

### C. elegans strains

*C. elegans* strains were maintained at 20 °C on nematode growth medium (NGM) plates previously seeded with OP50 *E. coli*. Strains harboring NMY-2 point mutations (Y163F and Y163E) were generated by CRISPR/Cas9. Briefly, the gonads of young adult *wild-type* animals (N2 strain) were injected with three different single guide RNA sequences (sgRNAs) and a single-stranded repair template (ssODN; IDT ultramer) to edit the *nmy-2* locus (details available in the supplementary material). The sgRNAs were individually expressed from the pDD162 vector, which also contains the Cas-9 gene re-encoded for *C. elegans* [93]. The injection mix also contained a sgRNA and a single-stranded repair template carrying the R92C mutation on the *dpy-10* gene, to allow the identification of successfully injected worms through the described roller phenotype [94, 95].The successful editing of the *nmy-2* locus was confirmed by genomic DNA sequencing. nmy-2 mutant strains were outcrossed six times with N2 animals to eliminate possible off target mutations.

### Brood size and embryo viability in C. elegans

Synchronized L1 animals of N2 and GCP693 strains were grown in NGM plates seeded with OP-50 *E. coli* at 20 °C for 72 h (1-day adults). Adult animals were singled out onto fresh NGM plates and the total number of eggs they laid over 72 h was counted for brood size quantification. For quantification of embryonic viability, the percentage of embryos that hatched was determined 24 h after embryos were laid.

### Cry5B intoxication assay

Synchronized L1 animals of N2 and GCP693 strains were grown in OP-50 *E. coli*-seeded NGM plates at 20 °C for 50 h (L4 stage) and transferred, every 2 days for a period of 5 days, to NGM plates previously seeded with control (empty vector pQE) or Cry5B-expressing BL21(*DE3*) *E. coli* [96]. Briefly, saturated cultures of *E. coli* transformed with the pQE empty vector or pQE-Cry5B vector were diluted (10x) in LB with 100 μ shaking. Expression of Cry5B was induced with 1 mM IPTG for 3 h at 30 °C. 100 µl of each culture at OD= 2±0.1 were plated onto NGM plates containing 2 mM IPTG and 100 μ ampicillin, and incubated at 25 °C overnight. Animals not responsive to touch were considered dead.

### Heat Shock assay

Synchronized L1 animals of N2 and GCP693 strains were grown as described above for 50 h. L4 animals of each strain were transferred to fresh NGM plates and maintained at 20 °C or incubated in a water bath at 35 °C for 1 h. Plates were then kept at 20 °C for 24 h. Surviving animals were singled out on fresh NGM plates and allowed to lay eggs for 24 h at 20 °C. Brood size and embryo viability were assessed 24 h later.

### Statistical analysis

Statistical analyses were carried out with Prism 8 (version 8.1.1), GraphPad Software, La Jolla California USA, www.graphpad.com), using one-way ANOVA with *Dunnett’*s *post hoc* analysis to compare different means in relation to a control sample and Tukey’s *post hoc* analysis for pairwise comparisons of more than two different means. Two-way ANOVA with Šídák’s *post hoc* analysis was used to compare each sample mean with another sample mean, in the same condition. Two-tailed unpaired and paired Student’s *t*-test was used for comparison of means between two samples.

## Funding

This work was funded by FEDER—Fundo Europeu de Desenvolvimento Regional funds through the COMPETE 2020—Operational Programme for Competitiveness and Internationalization (POCI), Portugal 2020, by Portuguese funds through FCT - Fundação para a Ciência e a Tecnologia/Ministério da Ciência, Tecnologia e Ensino Superior in the framework of the project POCI-01-0145-FEDER-030863 (PTDC/BIA-CEL/30863/2017) and by the European Research Council under the European Union’s Horizon 2020 Research and Innovation Programme (grant agreement 640553 – ACTOMYO - to AXC). CB, FSM and JMP were supported by FCT fellowships (SFRH/BD/112217/2015, SFRH/BPD/94458/2013 and SFRH/BD/143940/2019 respectively). CB was a Fulbright and FLAD fellow. SS and AXC receive support from the FCT Institutional and Individual CEEC program (CEECINST/00091/2018/CP1500/CT0006 and CEECIND/01967/2017, respectively).

## Acknowledgments

We are grateful to the members of i3S scientific facilities (ALM and Tracy) for technical assistance with microscopy and flow cytometry analysis, respectively; specially Maria Azevedo and Mafalda Sousa. We acknowledge the NHLBI Electron Microscopy Core Facility for the assistance during TEM samples processing. We thank Raffi Aroian (UMass Medical School) for sharing plasmids encoding Cry5b and respective control, Charles Bond (University of Pennsylvania) for helping with the analysis of motility assays and Renato Socodato (Glial Cell Biology, i3S) for initial assistance with FRET experiments.

## Author Contributions

Conceptualization: CB, FSM, DC and SS; Investigation, CB, FSM, DSO, JMP and NB; Funding Acquisition: JRS; AXC, DC and SS; Writing–Original Draft: CB and SS; Writing–Review & Editing: CB, FSM, DSO, JMP, NB, JRS; AXC, DC and SS.

## Abbreviations

ATP: Adenosine triphosphate

CA: Constitutively active

CFP: Cyan fluorescent protein

CTR: Control

DAPI: 4′,6-diamidino-2-phenylindole dihydrochloride

Dasa: Dasatinib

DMSO: Dimethyl Sulfoxide

DTT: Dithiothreitol

ELC: Essential light chain

EGTA: Egtazic acid

FAs: Focal adhesions

FRAP: Fluorescence recovery after photobleaching

FRET: Fluorescence resonance energy transfer

HMM: Heavy meromyosin

IPTG: Isopropyl-ß-D-thiogalactopyranoside

KD: Kinase dead

LLO: Listeriolysin O

MFI: Mean fluorescence intensity

MLCK: Myosin light chain kinase

MOI: Multiplicity of infection

MOPS: 3-(N-morpholino)propanesulfonic acid

NGM: Nematode growth medium

NI: Non-intoxicated

NMII: Non-muscle myosin II

NM2A: Non-muscle myosin 2A

NMHC2A: Non-muscle heavy chain myosin 2A

PFT: Pore-forming toxin

PI: Propidium Iodide

PM: Plasma membrane

PMSF: Phenylmethylsulfonyl Fluoride

pTyr: Tyrosine phosphorylation or phosphorylated tyrosine

RLC: Regulatory light chain

RNAi: Ribonucleic acid interference

WCL: Whole cell lysate

## Supplementary figures

**Figure S1. NMHC2A Tyr^158^ localization and conservation throughout evolution. (A)** Ribbon representation of the head domain of NMHC2A (green) to show Tyr^158^ (purple) and ADP (magenta) localization. The ELC (light yellow) and the actin-binding pocket (violet) are shown. PDB entry 1BR4. **(B)** NMHC2A amino-acid sequence analysis from different species and focused on the region involving the Tyr^158^, adapted from [25].

**Figure S2. Purified NM2A-GFP-Flag^HMM^ molecules from baculovirus-infected Sf9 cells. (A**) SDS-polyacrylamide gel stained with Coomassie Blue showing the separation of the purified heavy mero-myosins (HMM), and the regulatory and essential light chains (RLC and ELC, respectively) for each NM2A-GFP-Flag^HMM^ variant. **(B)** Representative single molecule electron microscopy images showing either a NM2A-GFP-Flag^HMM^-WT, Y158F or Y158E molecule. Scale bar, 10 nm. **(C)** Electron microscopy images showing the bound (arrow heads) and unbound (arrows) status of all purified NM2A-GFP-Flag^HMM^ variants to actin filaments in the absence or presence of ATP. Scale bar, 100 nm.

**Figure S3. Transfection and expression levels of the different NMHC2A variants in HeLa cells. (A)** Percentage of transfected cells determined by the quantification of GFP positive cells by flow cytometry. Each dot corresponds to a single independent experiment. Values are the mean ± SEM (*n* = 3) **(B)** Expression levels of the different constructs shown as MFI (mean fluorescence intensity) of the GFP positive cells, determined by flow cytometry. Each dot corresponds to a single independent experiment. Values are the mean ± SEM (*n* = 3).

**Figure S4. Quantification of the anisotropic distribution of stress fibers in HeLa cells ectopically expressing GFP-NMHC2A-WT or Y158F.** Each dot corresponds to a single cell. Values are the mean ± SEM (*n* > 200);

**Figure S5. Comparison of myosin protein sequence in different species.** Protein sequence analysis of the NMHC2A from *H. sapiens, M. musculus* and *C. elegans*, focused in the region containing the Tyr^158^ in human NMHC2A. The position of the corresponding residue in the other organisms is indicated.

## Supplementary Material

**Table 1.**
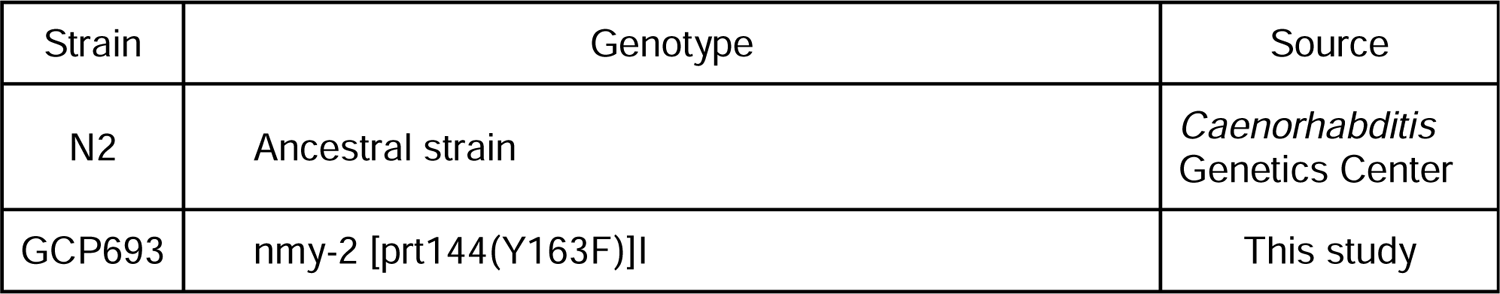
C. elegans strains used in this study.

**Table 2.**
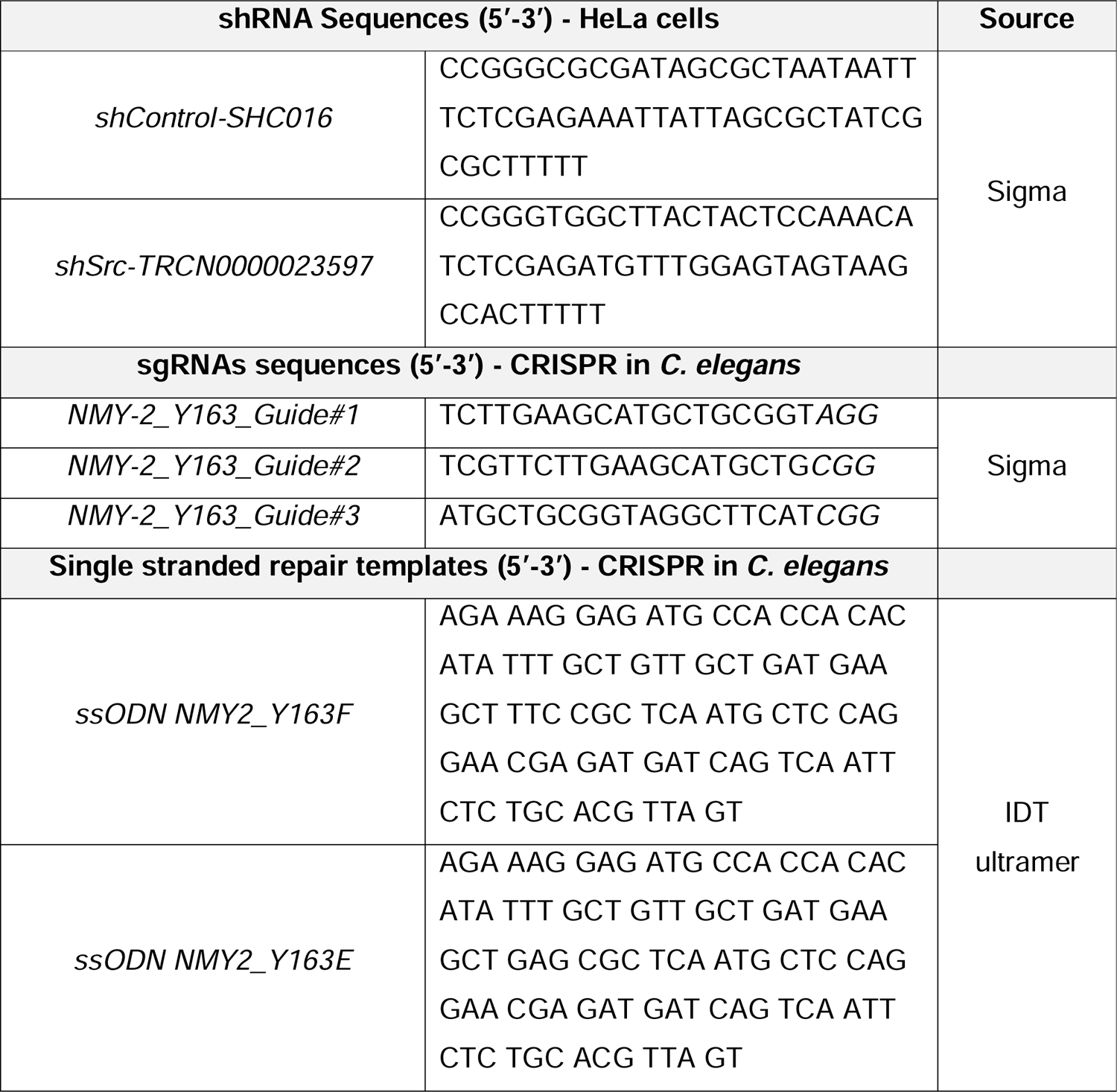
shRNA sequences for Src knockdown in HeLa cells and single-guide RNAs, single-stranded repair templates and RNAi sequences used in *C. elegans*

**Movie M1. Src is activated in response to LLO intoxication in HeLa cells.** HeLa cells expressing ECFP/YPet-based Src biosensor were imaged by time-lapse microscopy during 10 min (Non-intoxicated). LLO was added to a final concentration of 2 nM and cells were further imaged for 30 min. Sequential frames acquired every 30 s (15 frames per second display rate) show the ECFP (cyan) and YPet (yellow) fluorescence intensity separately, and the ECFP/YPet ratio. Scale bar, 10 μ

**Movie M2. NMHC2A pTyr does not affect the associated actin motility assay.** Rhodamine-phalloidin-labeled actin filaments moving on top of either the NM2A-GFP-Flag^HMM^ -WT, -Y158F or -Y158E. Sequential frames were acquired every 1 s for 2 min (15 frames per second display rate). Scale bar, 20 μ

**Movie M3. Ectopic expression of the different GFP-NMHC2A variants affects cell motility in HeLa cells.** HeLa cells ectopically expressing GFP-NMHC2A-WT or -Y158F were followed by time-lapse microscopy for more than 15 h. (Upper panel) Sequential frames acquired every 10 min (8 frames per second display rate). Overlay of the transmitted light and the fluorescence of GFP-NMHC2A variants is shown. Scale bar, 50 μm. (Lower panel) Representative tracking of the movement of cells expressing WT or Y158 versions of NMHC2A, produced by the Manual Tracking plug-in on Fiji™. Scale bar, 50 μm.

## Notes

### Competing Interest Statement

The authors have declared no competing interest.

