## Supplementary figures and images for "Src-dependent NM2A tyrosine-phosphorylation regulates actomyosin dynamics"

### Supplemental Figure S1

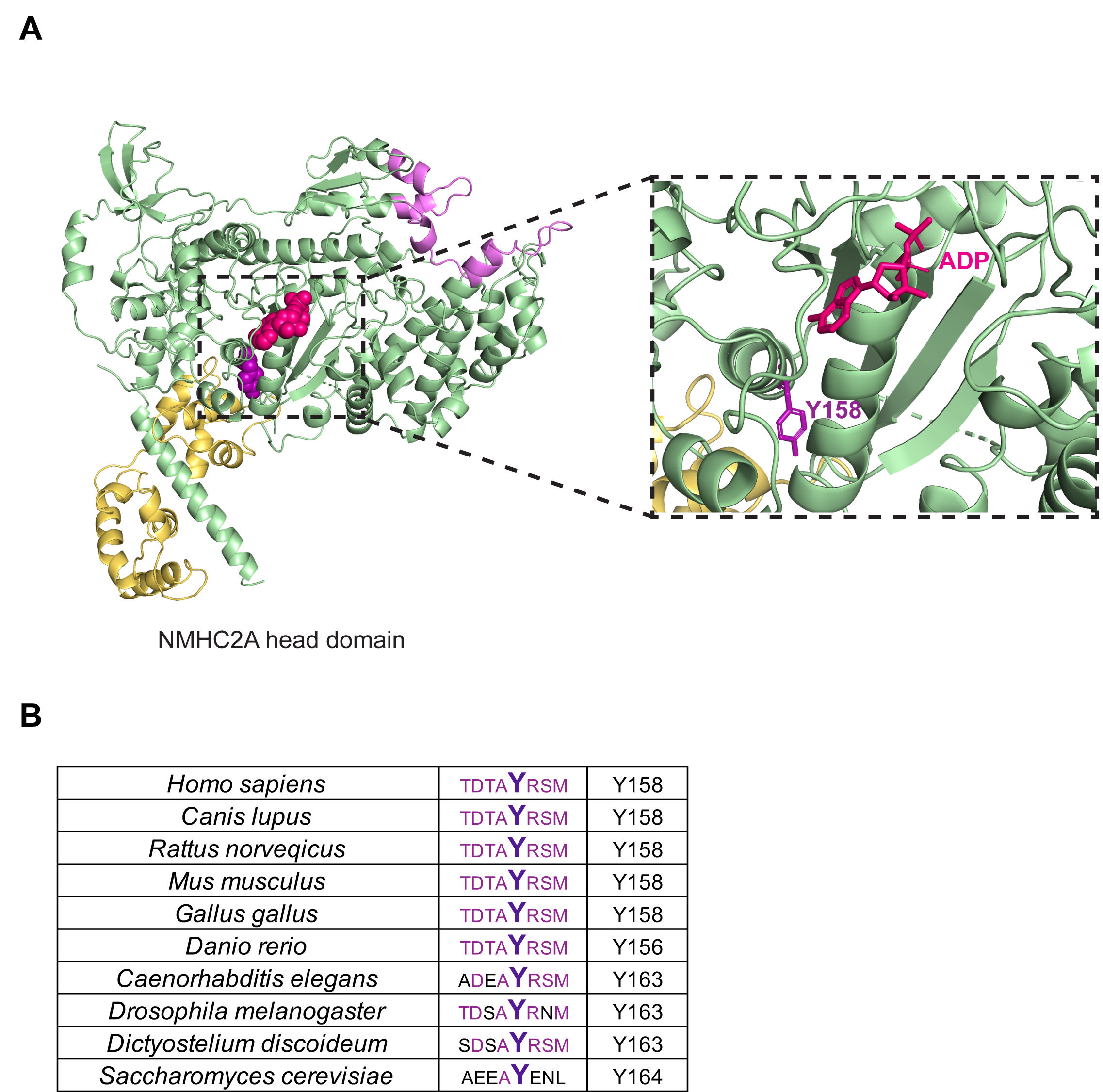

### Supplemental Figure S2

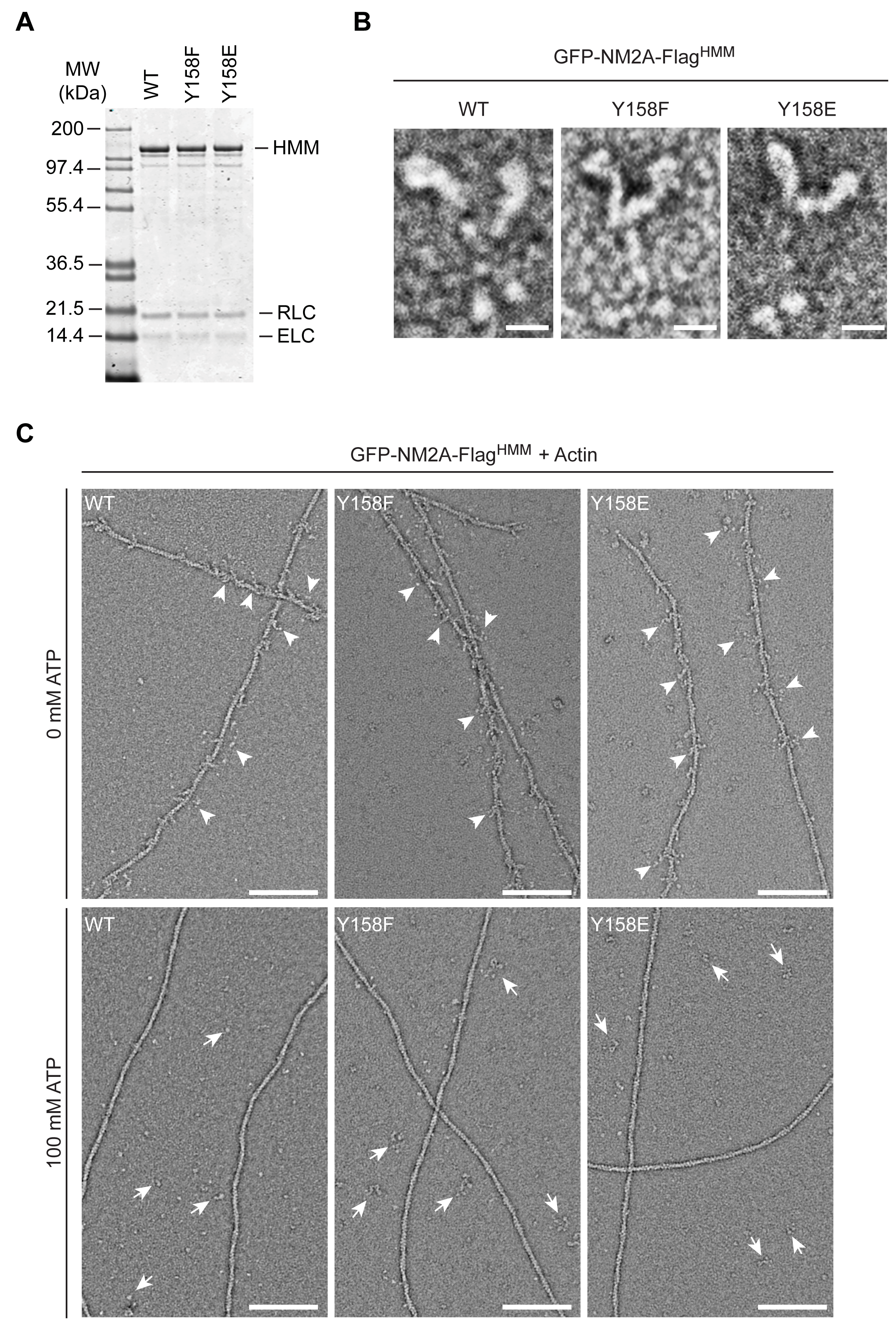

### Supplemental Figure S3

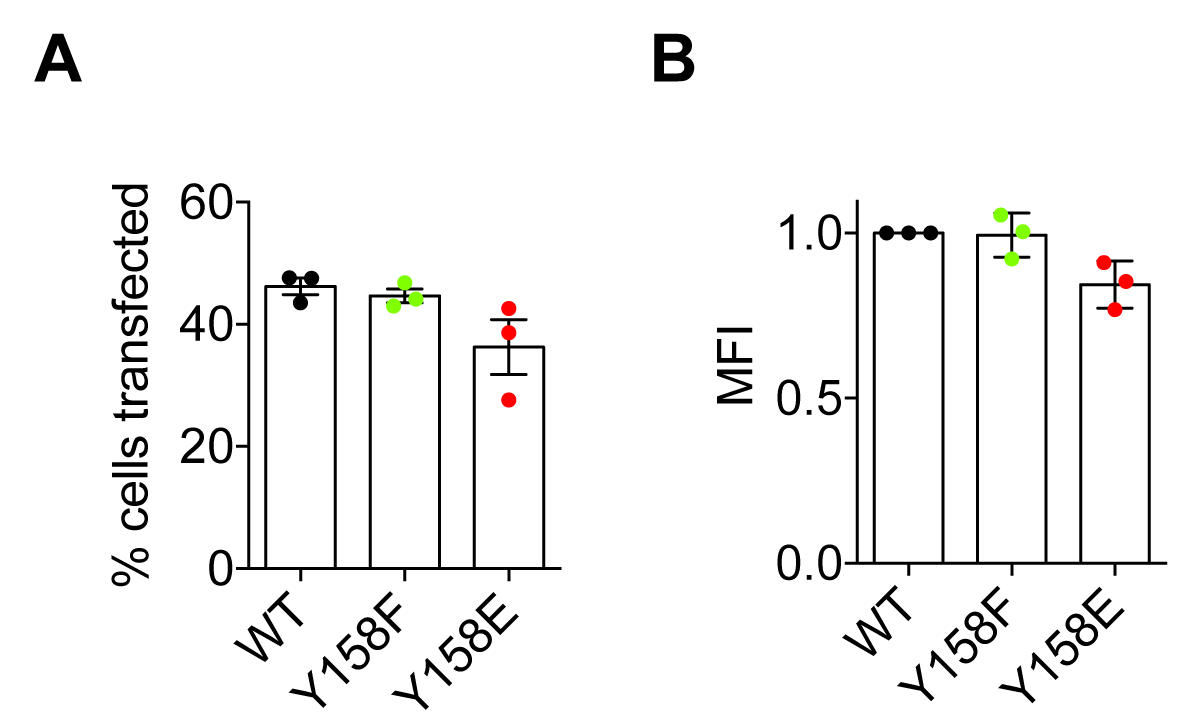

### Supplemental Figure S4

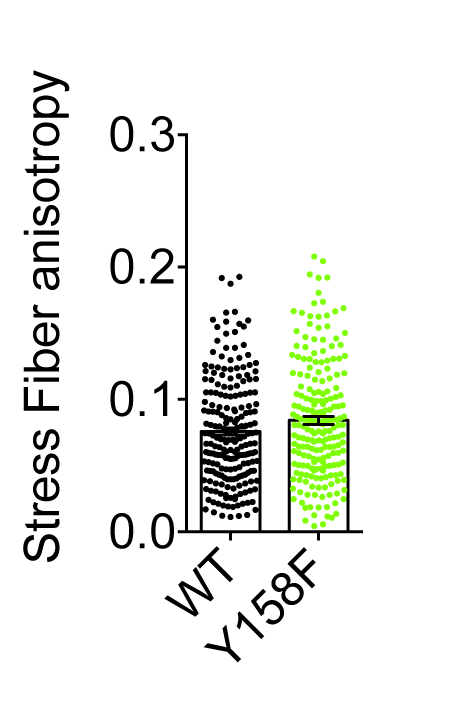

### Supplemental Figure S5

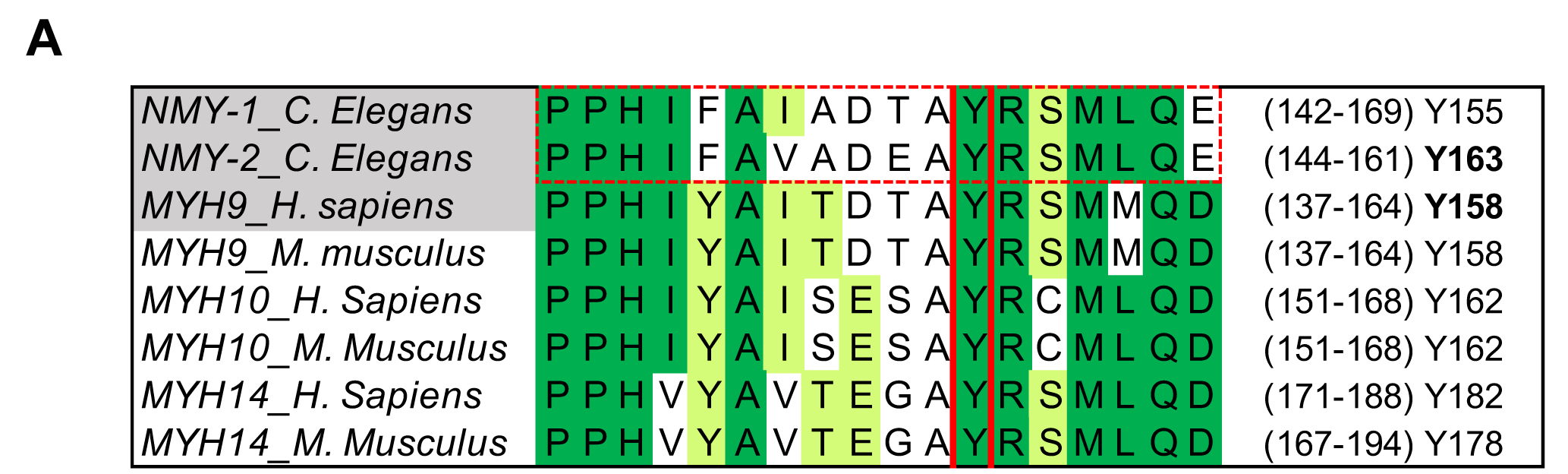
